# RNA-dependent association of the pyruvate dehydrogenase complex with mtDNA-containing assemblies supports mitochondrial translation

**DOI:** 10.64898/2026.08.26.747218

**Authors:** Johannes Hagen, Nupur Sharma, Felix Thoma, Veronika Iskra, Serena Schwenkert, Simon Schrott, Ignasi Forné, Julia Weisenseel, Mengqiao Yang, Nadja Lebedeva, Christof Osman

## Abstract

Mitochondrial DNA (mtDNA) encodes core subunits of the oxidative phosphorylation machinery, and its expression is spatially organized, with the genome packaged into nucleoids around which transcription and mitoribosome assembly are concentrated. How this architecture is built, and how it is linked to the metabolic state of the organelle, remains poorly understood. Here we show that the pyruvate dehydrogenase complex (PDHc), which supplies acetyl-CoA to the tricarboxylic acid cycle, is a component of this machinery in *Saccharomyces cerevisiae*. PDHc co-purifies with mtDNA and resides in high-molecular-weight assemblies whose integrity requires RNA rather than DNA, and proximity labeling places it selectively adjacent to the mitoribosome. The E1 subunit Pda1 concentrates into discrete foci that depend on mtDNA and disperse reversibly upon inhibition of mitochondrial translation. Loss of Pda1, Pdb1 or Lat1, but not of the E3-binding protein Pdx1, compromises the maintenance of mtDNA; deletion of *PDA1*, *PDB1* or *LAT1* also reduces output from a mitochondrially encoded reporter, and PDHc-deficient cells are hypersensitive to translational inhibition. Catalytically inactive Pda1 and Lat1 variants rescue both defects as effectively as the wild-type proteins, demonstrating a function genetically separable from acetyl-CoA synthesis. PDHc therefore acts as a non-catalytic component of the RNA-dependent architecture that supports mitochondrial gene expression.

## Introduction

Mitochondria are the principal sites of oxidative metabolism in eukaryotic cells, housing the tricarboxylic acid (TCA) cycle and the oxidative phosphorylation (OXPHOS) machinery. Although mitochondria retain their own genome, mitochondrial DNA (mtDNA) encodes only a small subset of OXPHOS subunits, and respiratory competence therefore depends on the coordinated expression of two genomes and on the faithful maintenance of mtDNA itself [1].

Within the matrix, mtDNA is packaged by dedicated HMG-box proteins — Abf2 in *Saccha-romyces cerevisiae* and TFAM in mammals — into nucleoids that organize the genome and serve as platforms for its replication and transcription [2–5]. The subsequent steps of gene expression are themselves spatially organized: in yeast, mitoribosomes reside in large assemblies that also contain transcription and RNA-processing factors [6], while in mammalian mitochondria, nascent transcripts are processed and matured within RNA granules — membraneless condensates that lie adjacent to nucleoids and serve as sites of mitoribosome assembly [7–10]. How this architecture is established, and how it is coordinated with the metabolic state of the organelle, remains poorly understood.

Notably, the nucleoid is not built from dedicated genetic factors alone. Alongside the packaging, replication and recombination machineries, unbiased crosslinking of proteins to yeast mtDNA by Butow and colleagues recovered several metabolic enzymes as nucleoid constituents [11], highlighting that nucleoid composition is remodeled in response to metabolic cues [12]. The best-characterised example is the TCA-cycle enzyme aconitase (Aco1), which binds mtDNA and contributes to its maintenance in a manner coupled to metabolic state [13, 14]. Intriguingly, subunits of the pyruvate dehydrogenase complex (PDHc) have likewise been found among nucleoid-associated proteins, both in yeast [14] and in vertebrate mitochondria [15], yet the basis and significance of this association have not been explored.

The PDHc catalyzes the oxidative decarboxylation of pyruvate to acetyl-CoA, providing a major entry point of glycolytic carbon into mitochondrial oxidative metabolism. In yeast, it comprises the E1 heterotetramer of Pda1 and Pdb1, a 60-mer core of the E2 dihydrolipoamide acetyltransferase Lat1, the E3 dihydrolipoamide dehydrogenase Lpd1, and the E3-binding protein Pdx1, assembling into a multivalent particle of roughly 9 MDa [16, 17]. Here we identify a non-catalytic role for the yeast PDHc in mitochondrial gene expression. We show that PDHc subunits associate with mtDNA-containing assemblies and are selectively proximal to the mitochondrial translation machinery, and that this association is mediated by RNA rather than by direct binding to the genome. Pda1 concentrates into discrete, mtDNA-dependent foci whose integrity requires ongoing mitochondrial translation. Loss of PDHc subunits compromises mtDNA maintenance and mitochondrial translation, and catalytically inactive variants rescue both defects, showing that these functions are genetically separable from acetyl-CoA production. Our results place a central carbon-metabolic enzyme within the RNA-dependent architecture of mitochondrial gene expression.

## Results

### Affinity purification of the mitochondrial nucleoid recovers the PDHc

As an exploratory approach to define the protein environment of the mitochondrial genome without prior assumptions, we affinity-purified the nucleoid via its principal packaging factor Abf2, tagged with a Spot tag. The tagged protein was expressed at levels comparable to untagged Abf2, and cells carrying Abf2–Spot as the sole copy of Abf2 showed none of the growth defects at 37 *^◦^*C characteristic of Δ*abf2* cells, indicating that the fusion is functional (Supplementary Fig. S1A, B). Purifications were performed from isolated mitochondria in parallel with untagged control strains, both with and without prior formaldehyde crosslinking. Enrichment was assessed by quantitative mass spectrometry as the intensity of each protein in the Abf2–Spot purification relative to the matched control, with the aim of identifying candidate nucleoid-associated proteins for subsequent targeted validation (Fig. 1A).

**Figure 1:**
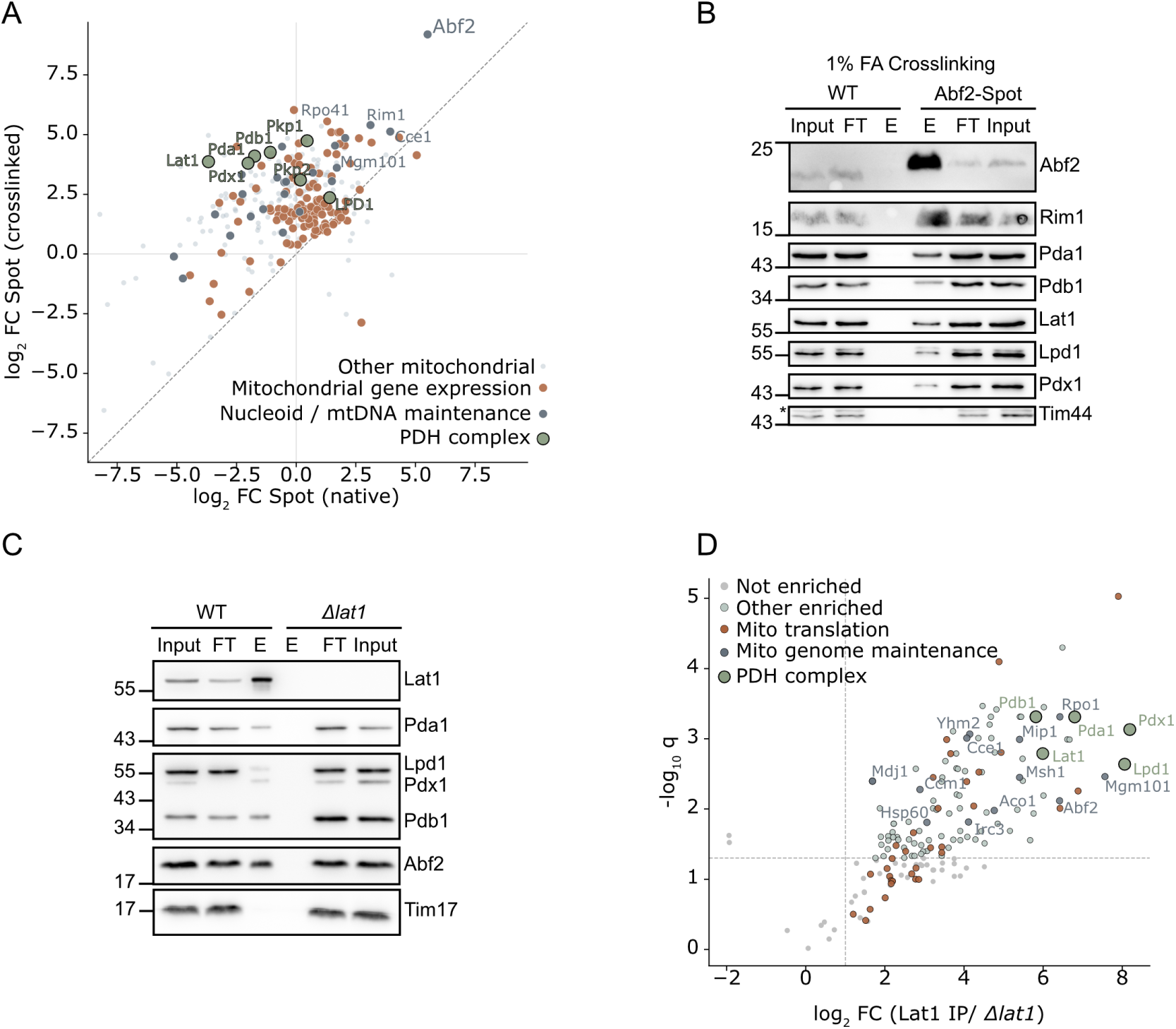
Association of PDHc subunits with mtDNA. **(A)** Quantitative mass spectrometry of Abf2–Spot affinity purifications versus untagged control, comparing log_2_ fold enrichment under native (*x*-axis) and 1% formaldehyde-crosslinked (*y*-axis) conditions (*n* = 1 exploratory purification; enrichment is reported as background-corrected log_2_ fold change without significance testing, see Methods). The dashed diagonal marks equal enrichment in both conditions. PDHc subunits (green), nucleoid / mtDNA-maintenance factors (dark grey), and mitochondrial gene-expression factors (orange) are highlighted; other mitochondrial proteins are shown in light grey. **(B)** Immunoblot analysis of Abf2–Spot affinity purifications following 1% formaldehyde crosslinking, comparing untagged wild-type (WT) and Abf2–Spot mitochondria. Input, flow-through (FT), and eluate (E) fractions were probed with the indicated antibodies (Tim44, control). Molecular masses (kDa) are indicated at left. **(C)** Immunoblot analysis of Lat1 immunoprecipitates from untagged wild-type (WT) and Δ*lat1* mitochondria. Input, flow-through (FT), and eluate (E) fractions were probed with the indicated antibodies (Tim17, control). Molecular masses (kDa) are indicated at left. **(D)** Volcano plot of Lat1 immunoprecipitate versus Δ*lat1* control (Welch’s *t*-test, Benjamini–Hochberg correction; *n* = 4 biological replicates). PDHc subunits (green) and factors involved in mitochondrial genome maintenance (dark grey) are highlighted; labeled proteins are those meeting the significance threshold (*q <* 0.05, log_2_ fold change *>* 1).

Consistent with specific isolation of the nucleoid, the Abf2 purifications recovered the established machinery of mitochondrial genome maintenance and expression. Enriched proteins included the mtDNA-packaging and maintenance factors Mgm101, Rim1, Cce1 and the mitochondrial RNA polymerase Rpo41, the mtDNA polymerase Mip1, and a broad set of mitochondrial gene-expression factors spanning the mitoribosome, mRNA-specific translational activators, mitochondrial RNA-processing enzymes and aminoacyl-tRNA synthetases (Fig. 1A, Supplementary Table S1). Recovery of these expected nucleoid- and expression-associated proteins confirmed that the approach captured *bona fide* constituents of the mitochondrial genome environment.

The Abf2 purifications also enriched the PDHc, but—unlike nucleoid maintenance and expression factors, which were enriched both natively and after formaldehyde treatment—PDHc subunits showed no enrichment under native conditions and became enriched only upon crosslinking (Fig. 1A), in line with earlier evidence for its association with mtDNA [14]. This crosslink-dependent enrichment encompassed all five subunits—the E1 components Pda1 and Pdb1, the E2 subunit Lat1, the E3 subunit Lpd1, and the E3-binding protein Pdx1—together with the regulatory kinases Pkp1 and Pkp2 and the related, E3-sharing 2-oxoglutarate dehydrogenase subunits Kgd1 and Kgd2. This behaviour indicates that the PDHc is spatially proximal to the nucleoid *in organello* but does not remain stably bound during native purification. The remaining enriched proteins comprised abundant mitochondrial matrix and respiratory-chain enzymes with no coherent functional signature beyond general metabolism, consistent with non-specific co-purification (Supplementary Table S1). To corroborate the mass-spectrometric result, we repeated the Abf2-Spot affinity purifications and probed for PDHc subunits by immunoblotting using newly generated antibodies, whose specificity was confirmed by loss of the corresponding signal in the respective deletion strains (Supplementary Fig. S4H). Co-purification of PDHc with Abf2–Spot was observed in three independent purifications, but relied on formaldehyde crosslinking (Fig. 1B, Supplementary Fig. S1C, representative blots), confirming the enrichment detected by mass spectrometry. Together, these data reveal an unexpected, crosslink-dependent association of the PDHc with the mitochondrial nucleoid.

As an independent, reciprocal test of this association, we performed affinity purifications from mitochondrial lysates using antibodies directed against the PDHc E2 subunit Lat1, with a Δ*lat1* strain processed in parallel as a specificity control. Immunoblot analysis of Lat1 immunoprecipitates revealed efficient recovery of Lat1 and co-purification of the remaining PDHc subunits Pda1, Pdb1, Lpd1, and Pdx1, confirming isolation of the PDHc (Fig. 1C). The mtDNA-packaging protein Abf2 was also specifically enriched in the Lat1 pulldowns, consistent with co-purification of mtDNA-associated material. Of note, in contrast to the Abf2-Spot purifications, no crosslinking was required to recover Abf2 together with Lat1.

To characterise the Lat1-associated proteome comprehensively, we analysed the immunoprecipitates by quantitative mass spectrometry and ranked interactors by the product of their enrichment and confidence (*π*-score; log_2_ fold change *× −*log_10_ *q*) [18]. All five PDHc subunits were among the most strongly enriched proteins, confirming the specificity of the purification (Fig. 1D). Of the thirty most enriched proteins, twenty-four were high-confidence mitochondrial proteins [19]. Beyond the PDHc itself, the most enriched mitochondrial proteins were dominated by factors of the mtDNA maintenance and expression machinery. These included the nucleoid and genome-maintenance components Abf2, Mgm101, Mnp1 and Sls1, the MutS homolog Msh1, the mitochondrial DNA polymerase Mip1 and the mitochondrial RNA polymerase Rpo41, together with components of the mitochondrial translation apparatus, including large- and small-subunit mitoribosomal proteins (e.g. Mrpl9, Mrpl11, Mrp51) and the translational activators Pet54 and Mtf2. Alongside these factors, we recovered a smaller number of other abundant mitochondrial proteins, predominantly respiratory-chain subunits and metabolite carriers (Supplementary Table S1).

### The PDHc subunit Pda1 is proximal to the mitochondrial ribosome *in vivo*

Biochemical co-purification cannot exclude association between mtDNA and PDHc subunits arises after lysis, and the Abf2 and Lat1 purifications had placed PDHc in the vicinity of both genome-maintenance factors and the translation machinery without resolving which of these it is physically adjacent to in living cells. We therefore turned to proximity-dependent biotinylation, which marks protein neighbourhoods *in vivo* within a limited labeling radius [20]. We fused the biotin ligase TurboID to the PDHc E1 subunit Pda1 and, as controls, expressed TurboID fused to a mitochondrial targeting sequence alone — a freely diffusing matrix enzyme (“Floaty–TurboID”) reporting nonspecific labeling — together with an untagged wild-type strain. Pda1–TurboID cells grew better than Δ*pda1* cells on non-fermentable carbon source (YPG) at 37 *^◦^*C, indicating that the fusion retains substantial Pda1 function (Supplementary Fig. S2A); immunoblotting confirmed expression of Pda1, albeit somewhat reduced levels relative to untagged Pda1 (Supplementary Fig. S2B).

Cells expressing Pda1–TurboID or Floaty–TurboID were grown in galactose-containing medium, and supplemented with Biotin for 30 minutes before mitochondrial purification, following which biotinylated proteins were affinity-purified on streptavidin beads (Fig. 2A). Immunoblotting revealed efficient biotinylation of the PDHc subunits Pda1, Pdb1, Lat1 and Pdx1 in Pda1–TurboID samples, whereas Lpd1 was not detectably biotinylated, potentially reflecting limited accessibility within the labeling radius (Fig. 2B). The mtDNA-packaging protein Abf2 was also biotinylated. By contrast, Floaty–TurboID and the untagged control showed essentially no capture of these proteins, confirming that labeling was both spatially specific and TurboID-dependent (Fig. 2B).

**Figure 2:**
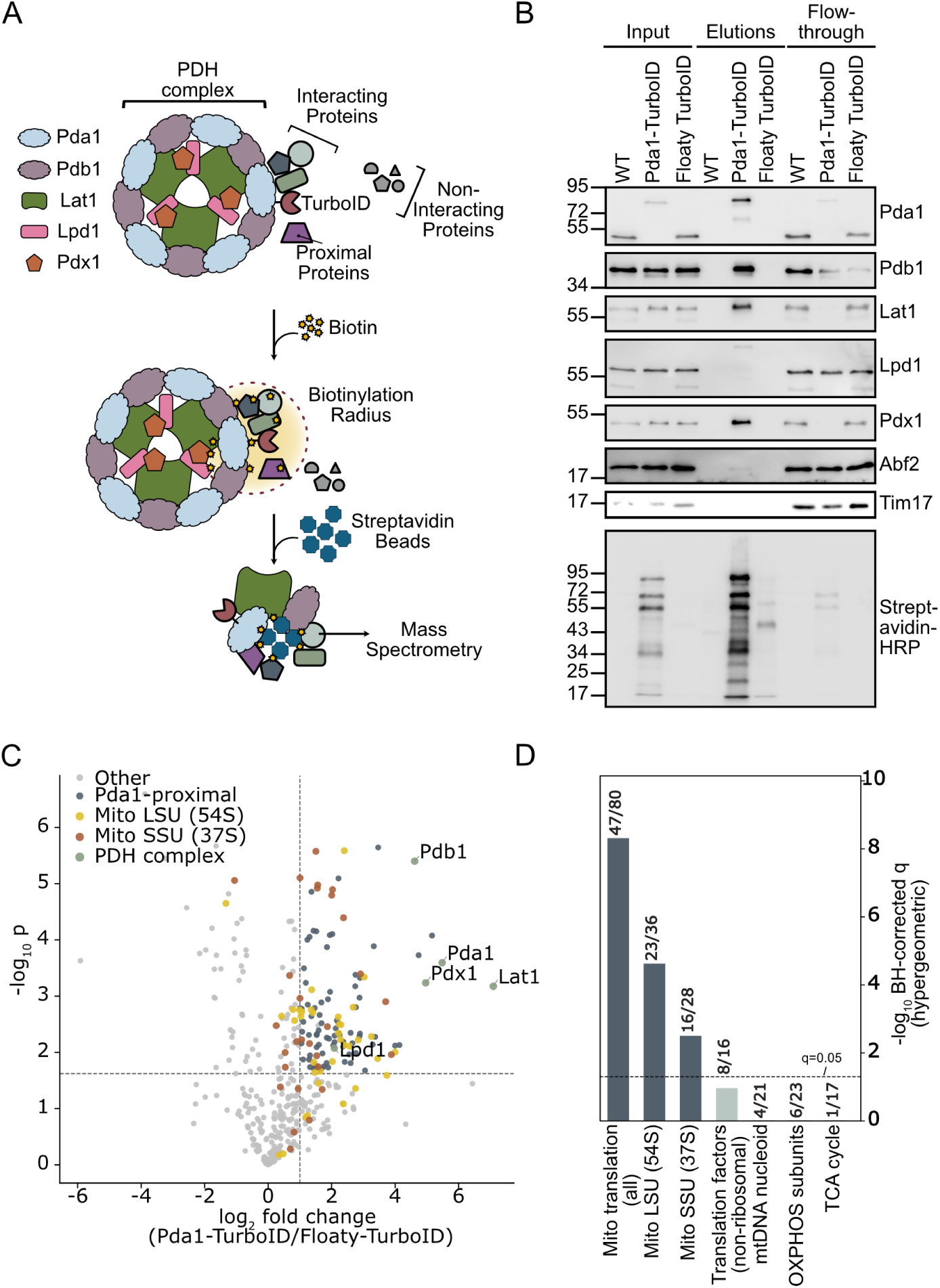
Proximity labeling using Pda1–TurboID. **(A)** Schematic illustration of the Pda1– TurboID proximity labeling strategy. **(B)** Western blot analysis of proteins labeled by proximity-dependent biotinylation in wild-type (WT), Pda1–TurboID and Floaty–TurboID cells. Biotinylated proteins were affinity-purified using streptavidin beads and detected by immunoblotting with the indicated antibodies (Tim17, control). Streptavidin blotting confirms overall biotinylation efficiency. **(C)** Volcano plot of streptavidin-purified proteins, Pda1–TurboID versus Floaty–TurboID (Welch’s *t*-test, Benjamini–Hochberg correction, *n* = 3 biological replicates). PDHc subunits (green), large ribosomal subunit proteins (54S, yellow), and small ribosomal subunit proteins (37S, orange) are highlighted; dashed lines indicate the significance and fold-change thresholds (*q <* 0.05, log_2_ fold change *>* 1). **(D)** Enrichment of curated mitochondrial protein sets among the Pda1-proximal proteome (hypergeometric test against the union of proteins detected across all conditions, Benjamini–Hochberg corrected). Bars show *−* log_10_ corrected *p*-values; numbers indicate proteins in the Pda1-proximal set relative to the total detected for each category. Dashed line, *q* = 0.05. Dark grey, significantly enriched; light grey, not significant; grey, comparison sets.

Comparative mass spectrometry of streptavidin-purified proteins showed that PDHc subunits were among the most strongly enriched proteins in the Pda1–TurboID samples relative to both the Floaty–TurboID (Fig. 2C) and the untagged control (Supplementary Fig. S2C). To ask which mtDNA-associated machinery Pda1 is proximal to, we tested curated gene sets — the large and small mitochondrial ribosomal subunits, non-ribosomal translation factors and aminoacyl-tRNA synthetases, and annotated nucleoid components — for over-representation among the proteins specifically enriched by Pda1–TurboID, with oxidative phosphorylation and TCA cycle enzymes included as controls for nonspecific labeling of abundant matrix proteins (Supplementary Table S1).

Among the tested categories, only the mitochondrial ribosome was significantly over-represented in the Pda1-proximal proteome, encompassing both the large (23 of 36 detected subunits; *q* = 2.4 *×* 10*^−^*^5^) and small (16 of 28; *q* = 3.1 *×* 10*^−^*^3^) subunits (Fig. 2D, Supplementary Table S1); as only proteins detected in the dataset entered the test, these denominators reflect the detectable subunits of each particle rather than its full complement. This enrichment was preserved when the analysis was restricted to proteins detected in all three Pda1–TurboID replicates, so that no foreground protein carried imputed values (*q* = 9.5 *×* 10*^−^*^4^ and 2.0 *×* 10*^−^*^3^), indicating that it is not an artifact of imputation. By contrast, non-ribosomal translation factors and aminoacyl-tRNA synthetases, oxidative phosphorylation subunits, TCA cycle enzymes and annotated nucleoid components were not enriched as categories, arguing against nonspecific labeling of abundant matrix proteins. Individual nucleoid factors were nonetheless biotinylated, most notably the mtDNA-packaging protein Abf2 (Fig. 2B). Thus, Pda1 is proximal to the mitochondrial ribosome while still contacting the mtDNA-packaging machinery, placing itself at a site where mtDNA-encoded OXPHOS subunits are synthesized.

### The PDHc resides in RNA-dependent high-molecular-weight mitochondrial assemblies

Although the proximity of the Pda1 to the mitoribosome pointed toward an RNA-mediated interaction, the physical coupling of mtDNA, nascent transcripts, and mitoribosomes meant that direct DNA binding could not be excluded *a priori*. To further examine the nature of this association, we tested whether it depends on RNA or DNA.

Mitochondria isolated from cultures grown in fermentable carbon sources (YPD, YPGal) or YPG were solubilised with digitonin and separated into supernatant and pellet fractions by centrifugation. Under standard conditions, 60–70% of each PDHc subunit (depending on the subunit) partitioned with the pellet fraction, despite PDHc being classically considered a soluble matrix enzyme. This pellet association was observed across all three carbon sources, indicating that it is not restricted to a particular metabolic state, although our fractionation is not quantitative enough to exclude carbon-source-dependent differences in the extent of association (Fig. 3A, Supplementary Fig. S3B,D,F,G,H). Treatment of the solubilised extract with DNase prior to centrifugation did not measurably alter this distribution (Supplementary Fig. S3F,G,H). In contrast, RNase treatment caused a pronounced redistribution of all PDHc subunits into the supernatant (Fig. 3A,B), regardless of the carbon source (Supplementary Fig. S3B,D). To confirm that both nucleases were active under these conditions, we verified in parallel control reactions that DNase and RNase efficiently degraded plasmid DNA and cellular RNA, respectively, when added to the digitonin-containing fractionation buffer (Supplementary Fig. S3A,C). These results indicate that the pellet association of PDHc is RNA-dependent and does not require intact mtDNA.

**Figure 3:**
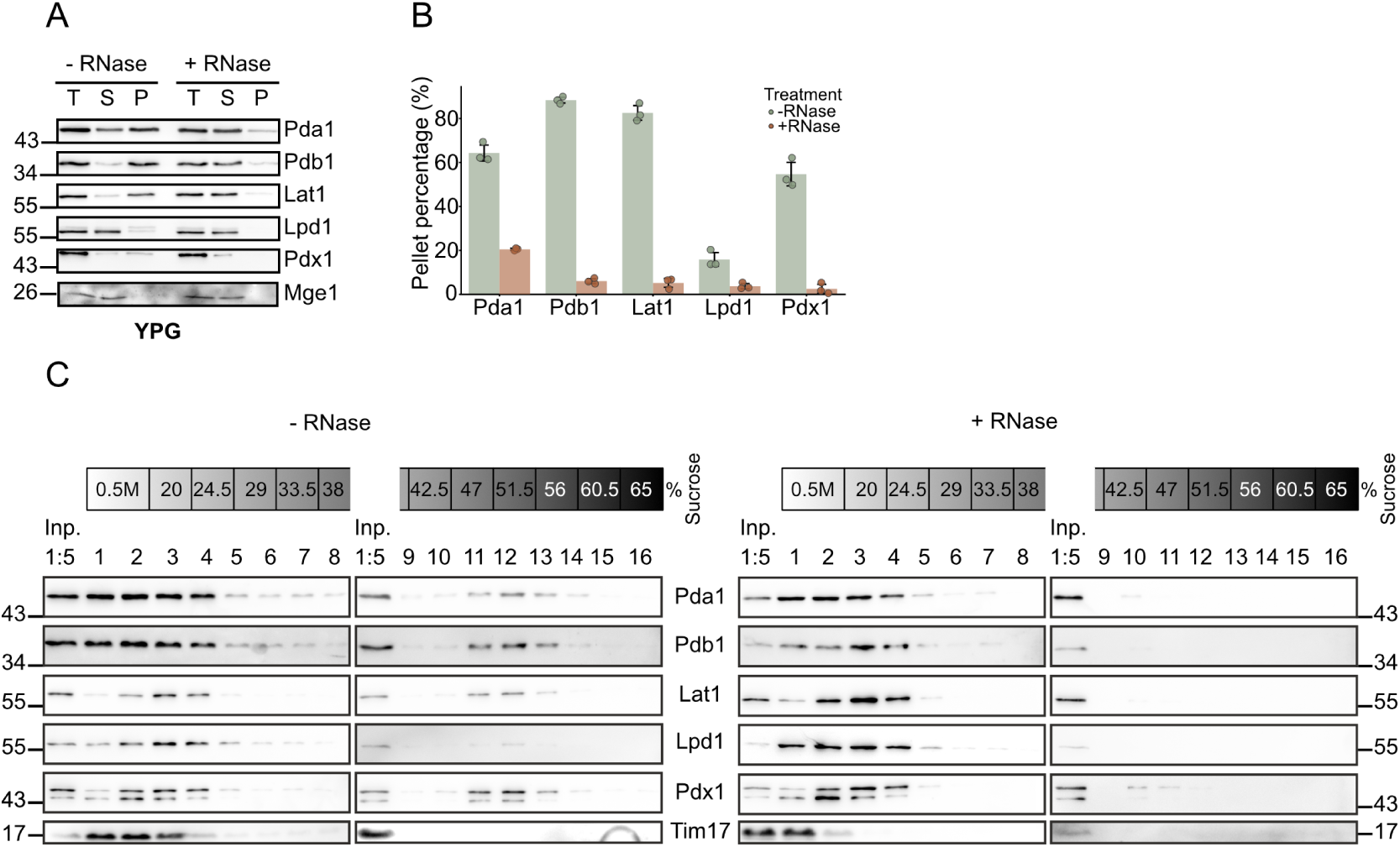
RNA-dependent association of PDHc with large macromolecular assemblies. **(A)** 1% Digitonin-solubilized mitochondria from YPG-grown cells were separated into total (T), supernatant (S), and pellet (P) fractions by centrifugation, with or without RNase treatment, and analyzed by immunoblotting with the indicated antibodies (Mge1, control). Molecular masses (kDa) are indicated at left. **(B)** Quantification of the proportion of each PDHc subunit in the pellet fraction from (A). **(C)** Digitonin-solubilized mitochondria were separated on 20–65% sucrose density gradients with or without RNase treatment, and fractions were analyzed by immunoblotting with the indicated antibodies. Input and gradient fractions 1–16 are shown (Tim17, control). Molecular masses (kDa) are indicated at the sides.

To corroborate this finding, we subjected digitonin-solubilised extracts to sucrose density gradient centrifugation (20–65% sucrose). In untreated samples, all PDHc subunits sedimented in heavy fractions, whereas RNase treatment shifted PDHc subunits almost entirely to light fractions (Fig. 3C). The same RNase-dependent shift was observed on YPD (Supplementary Fig. S3E). Together, the pellet fractionation and the gradient analysis converge on the same conclusion: a substantial fraction of the cellular PDHc pool resides in RNA-dependent high-molecular-weight mitochondrial assemblies. This finding supports the spatial proximity to the mitoribosome detected by Pda1–TurboID (Fig. 2A,C) and argues that PDHc is recruited to the mtDNA neighbourhood indirectly, via RNA-containing structures rather than via the genome itself.

### Pda1 defines discrete mtDNA-dependent assemblies within the mitochondrial network

To examine the spatial distribution of the PDHc within mitochondria, we next analyzed the localization of individual PDHc subunits by fluorescence microscopy. Each PDHc subunit was C-terminally tagged with mNeonGreen (NG), and functionality of the tagged proteins was assessed by growth analysis on YPG. Whereas deletions of *PDA1*, *PDB1*, *LAT1*, *LPD1*, or *PDX1* resulted in growth defects on YPG, particularly at elevated temperature (37 *^◦^*C), expression of the corresponding NG-tagged variants supported growth under these conditions, indicating that the tagged proteins are functional. The Lat1–NG fusion showed at most a minimal growth defect (Supplementary Fig. S4A). To assess whether the fluorescent signal reports the intact fusion proteins, we analysed each strain by immunoblotting with their respective antibodies and an anti-NG antibody. All fusions were detected at their predicted full-length molecular weight (Supplementary Fig. S4B). However, a faster-migrating species shifted by a constant *∼*23 kDa was also detected across the four subunits Pda1, Pdb1, Lat1 and Lpd1 (Supplementary Fig. S4C). Because this offset is very similar for the four unrelated proteins, it most parsimoniously reflects cleavage within the shared NG moiety rather than subunit-specific proteolysis, consistent with the documented backbone lability of fluorescent proteins near the chromophore [21, 22]. As the anti-NG epitope lies in the N-terminal portion of the tag, the detected fragment retains this epitope but has lost the C-terminal region that completes the chromophore, and the cleavage product is therefore expected to be non-fluorescent and not to contribute to the imaged signal. To visualize mitochondrial morphology, cells co-expressed a mitochondria-targeted red fluorescent marker (mKate2), and the distribution of the NG signal was analyzed relative to the mitochondrial network.

Strikingly, Pda1–NG exhibited a highly punctate distribution, typically forming fewer than five bright, discrete foci per cell together with several dimmer puncta along the mitochondrial network (Fig. 4A,B). Notably, Pda1–NG foci were readily detectable under both fermentative (YPD) and respiratory (YPG) growth conditions (Supplementary Fig. S4F), indicating that focal enrichment of Pda1 is not restricted to a specific metabolic state. In contrast, the remaining PDHc subunits displayed a largely diffuse mitochondrial distribution and did not form comparably prominent foci. This asymmetry was unexpected, since all five subunits belong to the same physical complex and would be expected to distribute together.

**Figure 4:**
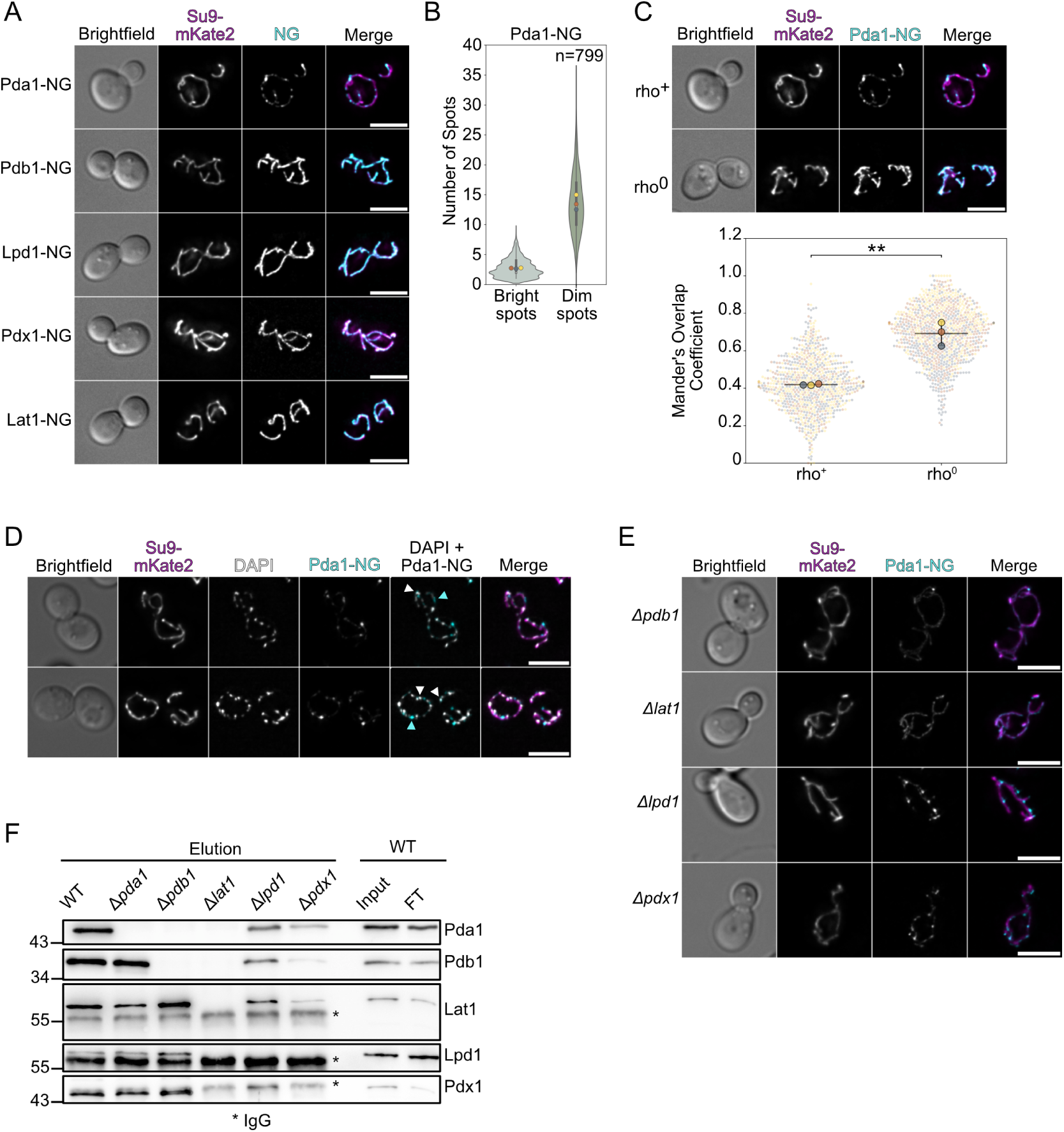
Pda1–NG assembles into foci within the mitochondrial network in an mtDNA-dependent manner. **(A)** Fluorescence microscopy of cells expressing the indicated PDHc subunit tagged with NG and the mitochondrial matrix marker Su9–mKate2. Scale bars, 5 µm. **(B)** Quantification of Pda1–NG spots per cell, classified as bright or dim. *n* = 799 cells from 3 biological replicates. **(C)** Fluorescence microscopy of Pda1–NG in mtDNA-containing (rho^+^) and mtDNA-lacking (rho^0^) cells (top), and Manders’ overlap coefficient between Pda1–NG and Su9–mKate2 (bottom). Scale bar 5µm. **(D)** Fluorescence microscopy of Pda1–NG cells co-stained with DAPI to visualize mtDNA. Cyan arrowheads, Pda1–NG foci lacking a coincident DAPI signal; white arrowheads, DAPI-stained nucleoids with an adjacent Pda1–NG focus. Two representative cells are shown; additional cells are shown in Supplementary Fig. S4E. Scale bars, 5 µm. **(E)** Fluorescence microscopy of Pda1–NG in Δ*pdb1*, Δ*lat1*, Δ*lpd1*, Δ*pdx1* **(F)** Immunoblot of Lat1 immunoprecipitates from the indicated strains (elution), with wild-type input and flow-through (FT) for reference, probed with antibodies against the indicated PDHc subunits. Asterisks mark IgG.

To exclude the possibility that the punctate localization of Pda1 reflects aggregation induced by the fluorescent tag—or an artifact of the NG cleavage described above—we generated an independent Pda1 fusion using the unrelated red fluorescent protein mScarlet. Pda1–mScarlet displayed a punctate localization pattern indistinguishable from that of Pda1–NG and was detected as a single full-length species without a comparable cleavage product (Supplementary Fig. S4D,G). Because NG and mScarlet are unrelated fluorescent proteins with distinct origins and biophysical properties, the observation of comparable foci with both tags argues against fluorophore-induced aggregation and confirms that the focal localization of Pda1 is independent of tag cleavage.

We next examined whether Pda1 foci depend on mitochondrial nucleic acids. In cells lacking mitochondrial DNA (rho^0^), the mitochondrial network appeared modestly altered, although the matrix marker mKate2 remained distributed throughout mitochondria (Fig. 4C). Strikingly, Pda1–NG no longer formed prominent discrete foci and instead exhibited a diffuse mitochondrial distribution. Quantitative analysis using Manders’ overlap coefficient revealed significantly increased overlap between the Pda1–NG signal and the mitochondrial marker in rho^0^ compared to rho^+^ cells, consistent with loss of focal enrichment in the absence of mtDNA. Together, these observations indicate that Pda1 puncta are not aggregation artifacts but instead depend on the presence of mitochondrial DNA and/or mtDNA-derived RNA, supporting the notion that Pda1 marks mtDNA-associated assemblies.

The mtDNA dependence of Pda1 foci raised the question of whether they coincide with the mitochondrial nucleoids themselves. In budding yeast, mtDNA is present in multiple copies packaged into nucleoids that are distributed as discrete foci along the mitochondrial network [23]. To address this, we counterstained rho^+^ cells expressing Pda1–NG with DAPI to visualise these nucleoids. Pda1 foci were found in the vicinity of DAPI-stained nucleoids but did not strictly coincide with them, with the two signals frequently offset within the mitochondrial network (Fig. 4D, Supplementary Fig. S4E). The lack of strict overlap is consistent with our biochemical finding that the association of PDHc with mtDNA-containing assemblies is RNA-rather than DNA-dependent, and supports a model in which Pda1 marks an mtDNA-associated compartment.

While only Pda1 appeared focally enriched, we tested whether the other PDHc subunits are required for its assembly into foci, imaging Pda1–NG in cells lacking individual partner subunits. In Δ*lat1* cells, Pda1 was entirely diffuse, whereas in Δ*pdb1* cells foci were largely lost: most cells displayed a diffuse Pda1 signal, with only occasional cells retaining a single focus. By contrast, foci were retained in Δ*pdx1* cells, indistinguishable from wild-type, and also in Δ*lpd1* cells, where they appeared, if anything, slightly more numerous. Thus, focus formation strictly requires Lat1 and largely depends on Pdb1, but not on the E3-binding protein Pdx1, and its persistence in Δ*lpd1* cells indicates that it does not require assembly of the complete holoenzyme (Fig. 4E).

To better understand these subunit requirements, we examined how the loss of each subunit affects the abundance and interactions of the others, by immunoblotting and by immunoprecipitation of Lat1 (Fig. 4F, Supplementary Fig. S4H,I). From wild-type mitochondria, Lat1 recovered all four partner subunits. The E3 module was dispensable for the integrity of the Pda1–Pdb1–Lat1 core: in Δ*lpd1* and Δ*pdx1* cells, Pda1 and Pdb1 remained associated with Lat1, and their steady-state levels were unchanged. As expected, Lpd1 was absent from Lat1 precipitates of Δ*pdx1* cells and Pdx1 from those of Δ*lpd1* cells. Interestingly, Pdb1 was required both for the stability and for the core association of Pda1: steady-state Pda1 was strongly reduced in Δ*pdb1* cells (Supplementary Fig. S4H), and the residual protein was no longer co-precipitated with Lat1. Pdb1 itself was still recovered with Lat1 in Δ*pda1* cells, indicating that it engages the Lat1 core independently of Pda1, whereas Pda1 associates with the core only when Pdb1 is present (Fig. 4F). Finally, Pda1 abundance was normal in Δ*lat1* cells, so the loss of foci in this background reflects a requirement for Lat1 in focus formation that is separate from Pda1 stability (Supplementary Fig. S4H). These subunit dependencies parallel the requirements for Pda1 foci, which persist where the Pda1–Pdb1–Lat1 core remains intact (Δ*pdx1*, Δ*lpd1*) and are lost or largely lost where it is disrupted (Δ*lat1*, Δ*pdb1*). It remains a conundrum, however, why Pdb1 and Lat1 do not themselves form comparable foci despite being required for them.

### The E1 and E2 PDHc subunits contribute to mitochondrial DNA stability independently of catalytic activity

To assess the functional consequences of disrupting the PDHc, we first examined the growth behaviour of strains lacking individual PDHc subunits. Deletions of *PDA1*, *PDB1*, *LAT1*, or *PDX1* resulted in only mild growth defects on YPD at both 30 *^◦^*C and 37 *^◦^*C (Fig. 5A). Notably, these strains also retained the ability to grow on YPG at 30 *^◦^*C, indicating that loss of PDHc activity does not preclude respiratory growth under standard conditions. Nevertheless, growth defects became apparent at 37 *^◦^*C. In contrast, deletion of *LPD1* caused a severe growth defect on YPG even at 30 *^◦^*C, consistent with Lpd1 serving as the shared E3 subunit of multiple mitochondrial dehydrogenase complexes, whose loss broadly compromises respiratory metabolism [24].

**Figure 5:**
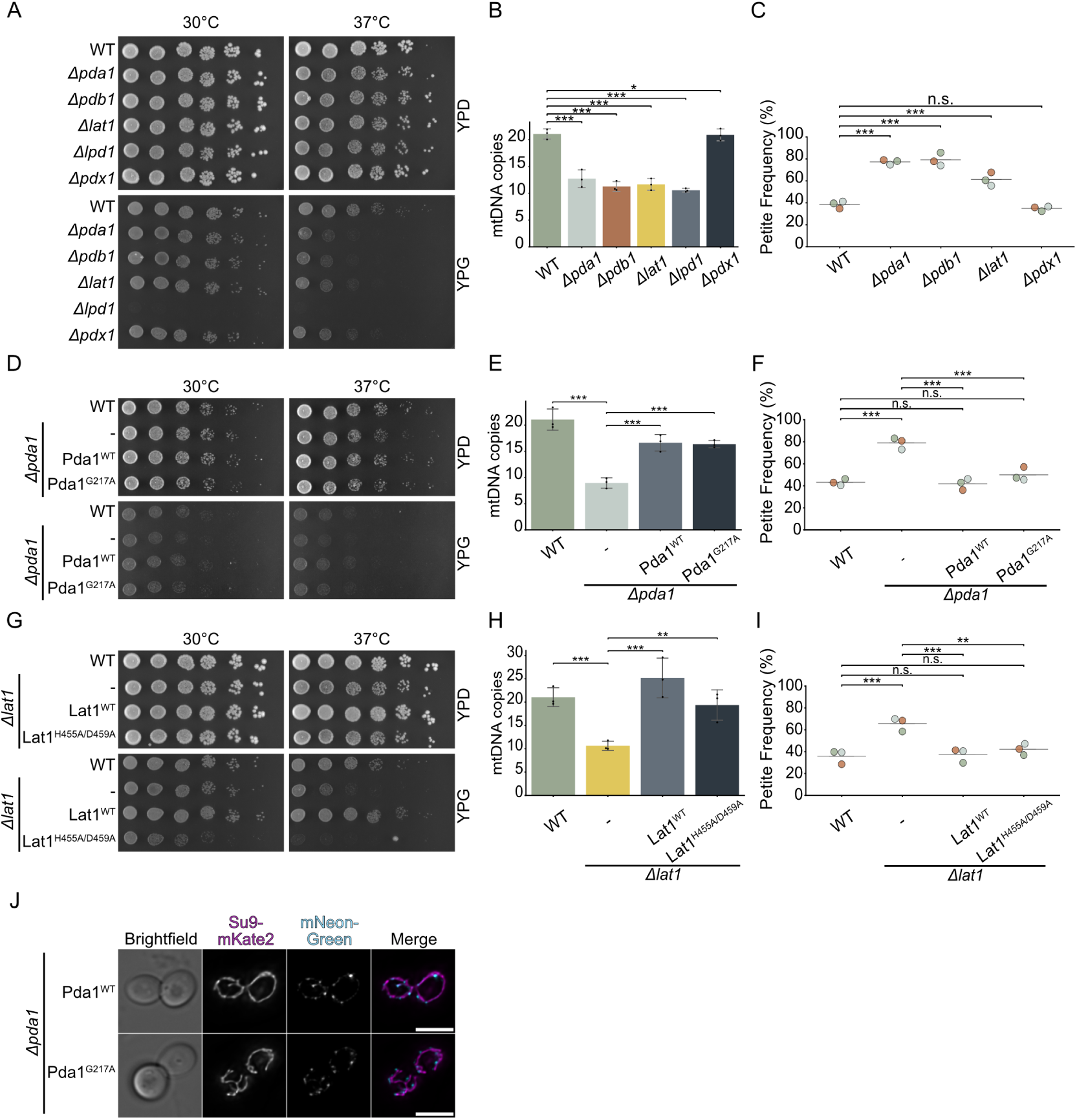
PDHc subunits fulfill a non-catalytic role in mitochondrial DNA maintenance. **(A)** Growth analysis of the indicated strains, on YPD and YPG medium at 30 *^◦^*C and 37 *^◦^*C. **(B)** Mitochondrial DNA (mtDNA) copy number of the indicated strains grown at 37 *^◦^*C, determined by qPCR (*COX1* /*ACT1*). *n* = 3 biological replicates. **(C)** Petite frequency of the indicated strains grown at 37 *^◦^*C. *n* = 3 biological replicates. **(D)** Growth analysis of Δ*pda1* cells expressing no PDHc variant (*−*), Pda1^WT^, or catalytically inactive Pda1^G217A^, on YPD and YPG at 30 and 37 *^◦^*C. **(E)** mtDNA copy number of the strains in (D) grown at 37 *^◦^*C determined by qPCR (*COX1* /*ACT1*). *n* = 3 biological replicates. **(F)** Petite frequency of the strains in (D) grown at 37 *^◦^*C. *n* = 3 biological replicates. **(G)** Growth analysis of Δ*lat1* cells expressing no PDHc variant (*−*), Lat1^WT^, or catalytically inactive Lat1^H455A/D459A^, on YPD and YPG at 30 and 37 *^◦^*C. **(H)** mtDNA copy number of the strains in (G) grown at 37 *^◦^*C determined by qPCR (*COX1* /*ACT1*). *n* = 3 biological replicates. **(I)** Petite frequency of the strains in (G) grown at 37 *^◦^*C. *n* = 3 biological replicates. **(J)** Fluorescence microscopy of Δ*pda1* cells expressing Pda1^WT^–NG or Pda1^G217A^–NG together with the mitochondrial matrix marker Su9–mKate2. Scale bars, 5 µm.

The mild phenotypes of most PDHc subunit deletions can be explained by alternative mitochondrial routes to acetyl-CoA [25, 26], most prominently the Ach1-dependent pathway, which generates acetyl-CoA from acetate [27] and becomes essential only when PDHc function is absent. Consistent with this, combining *ACH1* deletion with deletion of *PDA1*, *PDB1*, *LAT1*, or *PDX1* caused severe growth defects on YPG, demonstrating functional redundancy between the two pathways. As expected, *LPD1* deletion was epistatic to *ACH1*, as Δ*lpd1* cells already display a strong respiratory defect (Supplementary Fig. S5A).

We next asked whether PDHc subunits contribute to the maintenance of the mitochondrial genome itself. We therefore quantified mtDNA copy number in strains lacking individual subunits. At 30 *^◦^*C, copy number was little changed (Supplementary Fig. S5I); at 37 *^◦^*C, however, Δ*pda1*, Δ*pdb1*, Δ*lat1*, and Δ*lpd1* each showed a significant reduction, to approximately 50% of wild-type levels, whereas Δ*pdx1* showed no reduction at either temperature (Fig. 5B). The reduction in Δ*lpd1* cells should be interpreted with caution: because Lpd1 is shared among several mitochondrial dehydrogenase complexes and its loss abolishes respiration, its effect on mtDNA copy number is presumably indirect and pleiotropic rather than a specific consequence of disrupting PDHc.

To further assess mtDNA instability, we measured petite formation (Fig. 5C), excluding Δ*lpd1*, whose constitutive respiratory deficiency precludes meaningful petite scoring. In line with previous analyses, deletion of *PDA1*, *PDB1* [14], or *LAT1* produced a pronounced increase in petite frequency, apparent already at 30 *^◦^*C (Supplementary Fig. S5H) and marked at 37 *^◦^*C: whereas wild-type cells showed a petite frequency of approximately 40% at elevated temperature, Δ*pda1* cells reached up to 80% and Δ*lat1* cells exceeded 60%. In contrast, Δ*pdx1* showed no elevated petite frequency at any temperature, mirroring its unchanged mtDNA copy number.

The behaviour of Δ*pdx1* is notable in two respects. First, Pdx1 functions as the E3-binding protein that tethers the dihydrolipoamide dehydrogenase Lpd1 to the complex [28]; its dispensability for mtDNA maintenance therefore points to the Pda1–Pdb1–Lat1 core, rather than the fully assembled complex, as the relevant entity. This converges with our earlier observations that Pda1 foci persist in Δ*pdx1* cells and that the Pda1–Pdb1–Lat1 core remains intact in the absence of Pdx1 (Fig. 4E, F): the same three subunits — Pda1, Pdb1 and Lat1 — are required for focus formation and for genome maintenance, whereas the E3-binding protein is dispensable for both. Second, because Pdx1 is nonetheless required for wild-type PDHc catalytic output (Supplementary Fig. S5A), the absence of an mtDNA phenotype in Δ*pdx1* hints that catalytic output may not be the relevant determinant —a question we address directly below.

We considered two ways in which metabolism might still underlie the instability of mtDNA: altered pyruvate handling upstream of the complex, or residual catalytic activity of the remaining subunits. We addressed each in turn. To ask whether altered mitochondrial pyruvate utilization alone could account for the instability, we examined cells lacking Mpc1, the subunit common to both isoforms of the mitochondrial pyruvate carrier and therefore required for mitochondrial pyruvate import [29, 30]. In contrast to Δ*pda1* and Δ*lat1* cells, Δ*mpc1* cells showed only a minor increase in petite frequency, indicating that impaired pyruvate import per se is not sufficient to phenocopy the instability caused by loss of PDHc subunits (Supplementary Fig. S5B-D). Of note, the two perturbations are not metabolically equivalent—in Δ*mpc1* cells pyruvate cannot enter mitochondria, whereas in PDHc mutants pyruvate is imported but not converted to acetyl-CoA—but the absence of a phenotype despite disrupted pyruvate handling argues against a simple metabolic origin.

To test the second possibility—that residual catalytic activity, rather than the presence of the protein itself, is required for unperturbed genome maintenance—we reintroduced either the wild-type or a catalytically inactive variant of Pda1 (Pda1^G217A^) or Lat1 (Lat1^H455A/D459A^) into the corresponding deletion background and compared the two side by side. The catalytically inactive variants were expressed at levels comparable to their wild-type counterparts, as confirmed by immunoblotting (Supplementary Fig. S5F,G). Both inactive variants failed to rescue mild respiratory growth defects of the corresponding deletion mutants, but restored mtDNA copy number and reduced petite formation as effectively as reintroduction of the wild-type protein (Fig. 5D-I). As an internal control confirming loss of catalysis, neither variant rescued the severe growth defect of the corresponding *pdh*–*ach1* double mutant, in which acetyl-CoA production is essential (Supplementary Fig. S5E). Furthermore, Pda1^G217A^–NG retained the ability to form discrete mtDNA-associated foci indistinguishable from those of the wild-type fusion (Fig. 5J), demonstrating that catalytic activity is dispensable not only for mtDNA maintenance but also for assembly of Pda1 into the foci. Together, these results establish that the contribution of PDHc subunits to mtDNA maintenance is genetically separable from their canonical function in acetyl-CoA production, and indicate that the complex fulfills an additional, non-catalytic role in mitochondrial genome stability.

### The mtDNA-maintenance function of PDHc is linked to mitochondrial translation

The results so far established that the E1 and E2 PDHc subunits contribute to mtDNA maintenance through a non-catalytic function, but did not indicate at which step of mitochondrial gene expression this function acts. A long-standing observation provides a clue: in yeast, maintenance of intact, rho^+^ mtDNA depends on ongoing mitochondrial translation, as mutations that block mitochondrial protein synthesis, and inhibitors such as chloramphenicol, provoke loss of the genome [31, 32]. Given that PDHc is selectively proximal to the mitoribosome (Fig. 2) and resides in RNA-dependent assemblies (Fig. 3), we reasoned that its non-catalytic function might act at the level of transcription or translation. To distinguish these possibilities, we first asked whether loss of PDHc alters the mitochondrial transcriptome, by RNA sequencing of Δ*pda1* and wild-type cells grown in YPD at 30*^◦^*C.

The transcriptional response to Δ*pda1* was small and specific (Fig. 6A): of the *∼*6,000 genes tested, only 67 changed significantly (adjusted *p <* 0.1), and mitochondrial proteins were over-represented among them (odds ratio 3.2; *p* = 6.7*×*10*^−^*^5^). Critically, the mitochondrial transcriptome itself was essentially unchanged: all mtDNA-encoded OXPHOS transcripts — *COX1*, *COX2*, *COX3*, *COB*, *ATP6*, *ATP8*and *ATP9* — were unaltered, and the sole significantly changed mitochondrial transcript was *VAR1*, which notably encodes the only mitochondrially synthesised protein of the mitoribosome (base mean 1,228; *p*_adj_ = 0.0098). Nuclear genes for mtDNA maintenance and nucleoid function (*ABF2*, *MGM101*, *MHR1*, *MIP1*, *RPO41*, *RIM1*) were likewise unchanged, indicating that the genome-maintenance defect is neither accompanied nor compensated by transcriptional adjustment of the maintenance machinery. The changes that did occur formed a coherent metabolic signature of the lost acetyl-CoA step: the arginine-biosynthesis pathway, whose committed step consumes acetyl-CoA, was coordinately induced (*ARG1*, *ARG3*, *ARG5,6*, *ARG7*, *ARG8*, *CPA1*, *CPA2*), with the catabolic gene *CAR2* downregulated, and alternative fates for pyruvate were upregulated (*ALT1*, pyruvate *→* alanine; *PYC1*, pyruvate *→* oxaloacetate; *MPC2*, pyruvate import) — the expected consequences of removing the principal route from pyruvate to mitochondrial acetyl-CoA. This response was not a canonical retrograde signature: the retrograde reporter *CIT2* was downregulated rather than induced, indicating specific metabolic rerouting rather than generic mitochondrial-dysfunction signalling (Supplementary Table S2). Together, these data show that Δ*pda1* cells mount the metabolic adjustments expected from loss of PDHc while leaving the mitochondrial transcriptome and the mtDNA-maintenance machinery transcriptionally intact, arguing against a defect in mitochondrial transcription or transcript stability and directing attention to translation.

**Figure 6:**
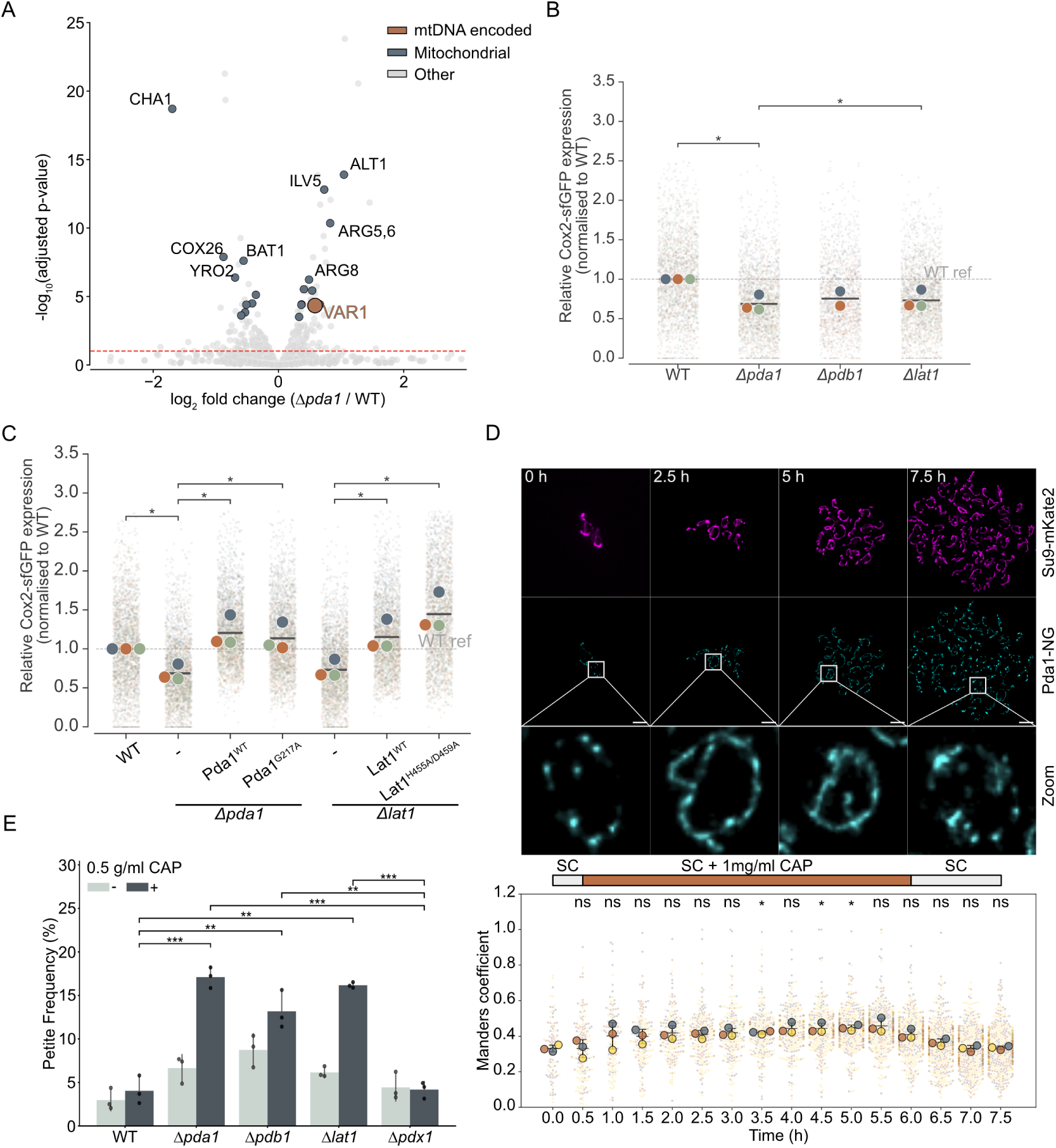
The mtDNA-maintenance function of PDHc is linked to mitochondrial translation. **(A)** Volcano plot of differential transcript abundance (Δ*pda1* versus wild-type) determined by RNA sequencing. mtDNA-encoded transcripts (orange) and nuclear-encoded mitochondrial transcripts (dark grey) are highlighted; all other transcripts are shown in light grey. The dashed line marks the significance threshold (adjusted *p* = 0.1). *n* = 3 (Δ*pda1*) and *n* = 2 (wild-type) biological replicates. **(B)** Cox2–sfGFP reporter expression in the indicated strains, measured by flow cytometry and normalized to wild-type (WT ref, dashed line). *n* = 3 biological replicates (for Δ*pdb1* n=2). **(C)** Cox2–sfGFP reporter expression in Δ*pda1* and Δ*lat1* cells expressing no variant (*−*), the wild-type protein (Pda1^WT^, Lat1^WT^), or the catalytically inactive variant (Pda1^G217A^, Lat1^H455A/D459A^), normalized to wild-type (WT). *n* = 3 biological replicates. **(D)** Live-cell imaging of Pda1–NG and the matrix marker Su9–mKate2 during chloramphenicol treatment and washout. Cells were imaged in synthetic medium (SC), during treatment with 1 mg/ml CAP, and after washout (SC), as indicated by the bar. Scale bars 5 µm. Bottom: Manders’ overlap coefficient between Pda1–NG and Su9–mKate2 over time (significance versus T_0_). *n* = 3 biological replicates. **(E)** Petite frequency of the indicated strains grown with or without 0.5 mg/ml chloramphenicol (CAP) at 30 *^◦^*C; *n* = 3 biological replicates.

We therefore examined mitochondrial translation using a reporter in which the *COX2* promoter drives expression of superfolder GFP (sfGFP) from within the mitochondrial genome [33], such that GFP fluorescence reflects the combined output of mitochondrial transcription and translation. Because this assay was performed on YPG, on which respiratory-deficient petite cells cannot proliferate, the growing population is strongly depleted of cells lacking functional mtDNA; any reduction in reporter output therefore most likely reflects a deficit within genome-competent cells rather than an increased proportion of petites. Deletion of *PDA1* or *LAT1* reduced steady-state sfGFP fluorescence at 30 *^◦^*C (Supplementary Fig. S6A), and this reduction was more pronounced at 37 *^◦^*C (Fig. 6B), paralleling the temperature dependence of the mtDNA-maintenance defect. Importantly, reintroduction of the catalytically inactive variants Pda1^G217A^ or Lat1^H455A/D459A^ restored reporter fluorescence as effectively as the wild-type proteins (Fig. 6C, Supplementary Fig. S6B), indicating that, as for mtDNA maintenance, PDHc subunits support mitochondrial translation through a non-catalytic function.

To test whether the Pda1 assemblies themselves are coupled to ongoing translation, we followed Pda1–NG foci in living cells during acute inhibition of mitochondrial translation. Addition of the mitoribosome inhibitor chloramphenicol led to progressive dispersal of Pda1 foci into a diffused mitochondrial distribution, while the mitochondrial network itself remained intact; upon washout of the drug, discrete foci reformed (Fig. 6D). The integrity of the Pda1 assemblies is thus reversibly dependent on active mitochondrial translation. Because chloramphenicol arrests elongating mitoribosomes without releasing them, the dispersal of the foci indicates that their integrity depends on ongoing translation elongation rather than on the mere presence of mitoribosomes.

This physical coupling was also mirrored physiologically: cells lacking PDHc subunits were hypersensitive to translational inhibition. Over five hours of chloramphenicol treatment at 30 *^◦^*C, wild-type cells showed only a marginal rise in petite frequency (from approximately 2.5% to 3%), whereas the same treatment sharply increased petite formation in Δ*pda1* and Δ*lat1* cells (from approximately 7% to 17%) (Fig. 6E). Consistent with every preceding assay, Δ*pdx1* behaved like wild-type and showed no increased sensitivity. A further, if indirect, line of evidence is genetic: a genome-wide genetic-interaction map [34] reported a negative interaction between *PDA1* and *MRS2*, which encodes a mitochondrial Mg^2+^ transporter required for group II intron splicing and efficient mitochondrial translation, and we confirmed this interaction directly (Supplementary Fig. S6C). Thus, whether mitochondrial translation is compromised pharmacologically or genetically, loss of the E1 and E2 PDHc subunits exacerbates mtDNA instability, revealing a functional interaction between PDHc and the translation machinery in the maintenance of the mitochondrial genome.

## Discussion

The PDHc is conventionally regarded as a soluble matrix enzyme dedicated to acetyl-CoA production. Our results reveal a second, non-catalytic role: in *S. cerevisiae*, a substantial fraction of PDHc resides in RNA-dependent, high-molecular-weight assemblies in the immediate neighbourhood of the mitochondrial gene-expression machinery, Pda1 concentrates into discrete mtDNA- and translation-dependent foci, and loss of PDHc subunits compromises mtDNA maintenance and mitochondrial translation through a function genetically separable from acetyl-CoA synthesis. PDHc thus joins a small group of metabolic enzymes that moonlight in the organization of the mitochondrial genome and its expression [13, 14, 35].

The genetic requirements for Pda1 foci and for mtDNA maintenance coincide: both depend on Pda1’s E1 partner Pdb1 and on the E2 subunit Lat1, but not on the E3-binding protein Pdx1 or the E3 subunit Lpd1, even though Pdx1 is required for full catalytic output. The relevant entity is therefore the Pda1–Pdb1–Lat1 core rather than the assembled holoenzyme. This is not contradicted by the Abf2 purifications recovering all five PDHc subunits: those pulldowns were performed in otherwise wild-type mitochondria upon crosslinking and report the complex as it normally exists at the nucleoid, whereas the deletion series identifies which subunits are functionally required. Physical proximity does not necessarily imply functional requirement, and the simplest reading is that the intact complex is recruited but that only the Pda1–Pdb1–Lat1 core is needed for mtDNA maintenance, with Lpd1 and Pdx1 dispensable for this role even when present. That Pdx1, required for full catalytic output, is not required here is consistent with the non-catalytic nature of the function; the same is predicted for the E3 subunit Lpd1, which is recruited to the PDHc by Pdx1. Its shared use by several 2-oxoacid dehydrogenase complexes and the glycine cleavage system, however, makes a Δ*lpd1* phenotype difficult to attribute to PDHc specifically. Why only Pda1 becomes focally concentrated, while the equally required Pdb1 and the far more abundant Lat1 remain diffuse, is not resolved by our data. The foci appear enriched in Pda1 rather than representing a miniature complex; Pdb1 and Lat1—and, by the same reasoning, Lpd1 and Pdx1—may be present below the level we can resolve, may act upstream to generate the recruited Pda1 species, or may reflect a distinct Pda1 assembly state. Quantitative two-colour imaging and higher-resolution approaches will be needed to define the composition and architecture of these structures.

A non-catalytic, nucleic-acid-associated role for a 2-oxoacid dehydrogenase subunit has precedents across life. Bacterial PDHc E2 binds regulatory DNA and influences transcription and replication initiation [36, 37], and the E1*β* and E2 subunits act in a sporulation checkpoint separable from catalytic activity [38]; the chloroplast PDHc-E2 subunit DLA2 binds the *psbA* mRNA and directs its translation [39, 40]; and the E2 of the related 2-oxoglutarate dehydrogenase is required for mitochondrial genome inheritance in *Trypanosoma brucei* [41]. Within mitochondria, the closest parallels are the moonlighting enzymes aconitase and Ilv5 [13, 14, 35] and, for the translation link, the fission-yeast aconitase Aco2, a fusion with a mitoribosomal protein essential for mitochondrial translation [42]. That PDHc can adopt such a spatial organization is itself not unprecedented. Metabolic enzymes readily assemble into higher-order structures whose formation is separable from catalysis: a survey of about 800 GFP-tagged yeast strains found that numerous enzymes of intermediary metabolism form punctate cytoplasmic foci in stationary phase, with the purine-biosynthetic enzyme Ade4 and glutamine synthetase Gln1 cycling reversibly between focal and diffuse states as adenine or glucose is withdrawn and restored [43, 44], and CTP synthase polymerises into filaments that can serve a structural role independent of catalysis [45]. Such assemblies also form within mitochondria: cryo-electron tomography of meiotic budding yeast resolved filaments of the matrix aldehyde dehydrogenase Ald4, alongside cytoplasmic polymers of acetyl-CoA synthetase Acs1 that are required for aged spores to resume growth [46]. Finally, a comparable spot-like organisation of PDHc itself has been described in mammalian cells, where a GFP-tagged E1*α* subunit — the orthologue of Pda1 — and the endogenous complex both form discrete, temporally stable puncta along the mitochondrial network [47]. In yeast, moreover, early biochemical fractionations of mitochondrial nucleoids reported the HMG-box protein Mnp1 together with nearly all PDHc subunits [48], an association whose functional significance had remained unexplored. A non-enzymatic role for yeast PDHc has also been reported in mitophagic trafficking, distinct from the function described here [49]. Our findings extend this pattern to the mitochondrial PDHc of budding yeast and, unlike the DNA-directed bacterial cases, identify the association as RNA-mediated.

Where within gene expression does PDHc act? Three observations place it downstream of transcription and at the mitoribosome: the association is RNA- but not DNA-dependent, the Pda1-proximal proteome is selectively enriched for mitoribosomal subunits, and mitochondrial transcripts are unchanged in Δ*pda1* cells while output from a mitochondrially encoded reporter is reduced. As a working model, we propose that the PDHc core is a constituent of the RNA-containing, mitoribosome-associated compartments in which mitochondrial transcripts and the mitoribosome are co-organized [6, 9, 50], and that it supports the integrity or organization of that compartment; because maintenance of rho^+^ mtDNA requires ongoing mitochondrial translation [31, 32], a translational deficit would in turn destabilize the genome. This ordering is supported by the reporter defect appearing already at 30 *^◦^*C, where mtDNA copy number is largely unchanged, by the hypersensitivity of PDHc-deficient cells to chloramphenicol, and by the genetic interaction between *PDA1* and *MRS2*, which is required for mitochondrial intron splicing and translation. That catalytically inactive PDHc restores both mtDNA maintenance and reporter expression excludes a mechanism based on locally generated acetyl-CoA. Our data do not distinguish whether PDHc contributes to mitoribosome assembly, to mRNA delivery, or to the physical coherence of this compartment, nor have we shown that any subunit binds RNA directly; defining the step and identifying the RNAs involved, for instance by CLIP-based approaches, are the immediate next questions.

Why a pyruvate-metabolizing enzyme should take on this role is unresolved, but its metabolic position is suggestive. PDHc catalyses the committed entry of carbon into mitochondrial oxidative metabolism, and, with the sole exception of the mitoribosomal protein Var1, every mtDNA-encoded protein is an OXPHOS subunit; embedding a role in OXPHOS-subunit synthesis within the entry enzyme would couple carbon flux to gene expression without a dedicated signalling relay — an arrangement made plausible by the reciprocal catalysis/RNA-binding switches of DLA2 and cytosolic aconitase [39, 51] and by the extensive phosphoregulation of PDHc [52]. This coupling remains hypothetical: we detect the association under both fermentative and respiratory conditions and have no evidence that it is itself metabolically regulated. Whether an equivalent function operates outside budding yeast remains to be tested, although a recent multiomic survey of mammalian cells with mitochondrial gene perturbations is suggestive: across more than 200 knockout lines, the abundance of PDHc subunits emerged among the strongest positive predictors of mtDNA copy number, alongside the packaging factor TFAM [53]. This correlation was not pursued in that study, but its direction matches the loss of mtDNA we observe in yeast PDHc mutants and suggests that the connection between PDHc and mitochondrial genome maintenance may be conserved.

A separate connection between PDHc and mitochondrial gene expression has been reported for human LETM1 (yeast homologues Mdm38 and YLH47), which associates with the mitoribosome and is required for mitochondrial translation and mtDNA organization, and which regulates PDHc activity; PDHc itself co-fractionates with these mitochondrial nucleoprotein complexes [54], linking the same elements — the mitoribosome, mtDNA maintenance, and PDHc — that our data connect in *S. cerevisiae*. Given that PDHc deficiency causes severe human disease attributed to defective pyruvate oxidation [55], it may be worth asking whether compromised mitochondrial gene expression contributes to the pathology.

## Materials and Methods

### Yeast strains, plasmids, and growth conditions

All experiments were performed in *Saccharomyces cerevisiae* derived from the W303 genetic background. A complete list of strains and plasmids used in this study is provided in Supplementary Table S3 (used oligonucleotides can be made available upon reuqest). Unless stated otherwise, cells were grown in rich medium containing 1% yeast extract, 2% peptone and 0.004% adenine (all w/v) supplemented with either 2% glucose (YPD), 2% galactose (YPGal) or 3% glycerol (YPG) as the carbon source. Cultures were maintained at 30 *^◦^*C; growth at elevated temperature was assessed at 37 *^◦^*C where indicated. Selection of transformants was performed on synthetic complete medium lacking the appropriate amino acid (0.67% (w/v) yeast nitrogen base, 0.192% (w/v) drop-out mix minus uracil (US Biological D9535), 2% (w/v) glucose), for synthetic complete selection plates 0.1% (w/v) 5-fluoroorotic acid monohydrate (5-FOA) and 0.005% (w/v) uracil was added to the SC-Ura plates or on rich medium containing the relevant antibiotic (300 µg/ml hygromycin B, 300 µg/ml G418, 100 µg/ml nourseothricin).

Gene deletions and C-terminal endogenous tagging were generated by PCR-based homologous recombination using pFA6a/pYM-based cassettes [56], conferring resistance to hygromycin B (*hphNT1*), nourseothricin (*natNT2*), or G418 (*kanMX4*). Where a prototrophic marker was required, a *URA3* cassette flanked by two *hisG* repeats was integrated at the *LEU2* locus and subsequently excised by counterselection on 5-fluoroorotic acid (5-FOA). Correct integration was verified by colony PCR and, for tagged strains, by immunoblot and sequencing. C-terminal fluorescent fusions were generated with mNeonGreen, mScarlet-I, or mKate2; matrix-targeted, non-fused fluorescent controls were generated by fusing mNeonGreen or mKate2 to the Su9 presequence (subunit 9 of the *Neurospora crassa* F_0_-ATPase). The biotin ligase TurboID was fused C-terminally to Pda1; a matrix-targeted, freely diffusing control (“Floaty–TurboID”) was generated by fusing TurboID to the Su9 presequence alone.

### Growth analysis

Logarithmically growing cells were adjusted to a starting concentration of 3.2 *×* 10^6^ cells/ml and serially diluted 1:5. From each dilution, 3.5 µl was spotted onto YPD or YPG plates, which were incubated for 24 - 48 h prior to imaging with a Vilber Fusion FX imaging system.

### Catalytically inactive PDHc variants

Catalytically inactive alleles of *PDA1* and *LAT1* were generated by site-directed mutagenesis of the corresponding active-site residues (*PDA1* ^[G217A]^ and *LAT1* ^[H455A/D459A]^) and expressed in the respective deletion background from *HO* (Lat1) or *LEU2* (Pda1) locus. Loss of catalytic activity was confirmed functionally by the failure of the variants to rescue the respiratory growth defect of the corresponding Δ*pdh*Δ*ach1* double mutant (Fig. 5).

### Lysis of yeast cells for immunoblot analysis

Yeast cells were grown in the desired medium or on plates, and 2.5 OD_600_ units of cells were harvested by centrifugation at 5,000 rpm for 5 min. The supernatant was discarded, and the pellet was resuspended in 100 µl H_2_O. After addition of 100 µl 0.2 M NaOH, samples were briefly vortexed and incubated for 5 min at room temperature. Cells were pelleted again by centrifugation at 5,000 rpm for 5 min, the supernatant was discarded, and the pellet was resuspended in 100 µl 1*×* SDS sample buffer (60 mM Tris-HCl pH 6.8, 5% glycerol, 2% SDS, 4% *β*-mercaptoethanol, 0.0025% bromophenol blue). Samples were incubated at 95 *^◦^*C for 3 min, centrifuged for 30 s at 13,000 rpm, and 6 µl of the resulting supernatant was loaded per lane for SDS-PAGE.

### Isolation of crude mitochondria

Mitochondria were isolated by differential centrifugation following spheroplast lysis. Cells were grown to mid-log phase (OD_600_ *≈* 0.8–1.2) in YPGal medium unless otherwise indicated, harvested, and washed. Cell walls were pre-treated with alkaline solution (100 mM Tris, 10 mM DTT; 10 min, 30 *^◦^*C), and cells were converted to spheroplasts by treatment with Zymolyase 20T (Amsbio, 120491-1; 6–6.6 mg/ml) for 30 min at 30 *^◦^*C in spheroplast buffer (20 mM KH_2_PO_4_ pH 7.4, 1.2 M sorbitol). Spheroplasts were resuspended in homogenization buffer (10 mM Tris-HCl pH 7.4, 0.6 M sorbitol, 1 mM EDTA, 0.2% BSA [fatty acid-free], 1 mM PMSF) and lysed by Dounce homogenization (15 up-and-down strokes, 100 ml Kontes Dounce homogenizer). Mitochondria were recovered by differential centrifugation (low-speed clearing: 3,000 rpm, 5 min, 4 *^◦^*C to remove unbroken cells and nuclei; mitochondrial pelleting: 10,000 rpm, 10 min, 4 *^◦^*C), followed by further purification through a sucrose cushion (SEM500 buffer: 500 mM sucrose, 1 mM EDTA, 10 mM MOPS-KOH pH 7.2; 13,000 rpm, 10 min, 4 *^◦^*C). Purified mitochondria were resuspended in SEM buffer (250 mM sucrose, 1 mM EDTA, 10 mM MOPS-KOH pH 7.2). Protein concentration was determined by Bradford assay. Mitochondria were either used fresh or snap-frozen in liquid nitrogen and stored at *−*80 *^◦^*C.

### Immunoprecipitation, proximity labeling, and mass spectrometry

**Abf2–Spot affinity purification** Affinity purification of Abf2-Spot was performed from isolated mitochondria (YPG grown). For crosslinked samples, 2 mg mitochondria were resuspended in 1% formaldehyde (in SEM buffer: 250mM Sucrose, 1 mM EDTA, 10 mM MOPS-KOH, pH7.2) and incubated for 10 min at room temperature; crosslinking was quenched with Tris-HCl pH 8.0 (750 mM final) for 1 min, and mitochondria were washed once in detergent-free lysis buffer to remove residual formaldehyde. Non-crosslinked samples were processed in parallel without fixation. Both were then solubilized in lysis buffer (15 mM Tris-HCl pH 7.4, 80 mM KCl, 2 mM EDTA pH 8.0, 0.2 mM spermine, 0.5 mM spermidine, EDTA-free protease inhibitor) supplemented with 0.5% NP-40 (2 mg mitochondria per pulldown) for 15 min on ice and cleared by centrifugation (1,000*× g*, 5 min, 4 *^◦^*C). Cleared lysate (2% retained as input) was incubated with Spot-Trap Magnetic Particles M-270 (ChromoTek; 5 *µ*l slurry per mg mitochondria) for 2 h at 4 *^◦^*C, and the unbound fraction was collected as flowthrough (2%). Beads were washed twice in wash buffer 1 (lysis buffer + 0.1% NP-40) and twice in wash buffer 2 (detergent-free lysis buffer).Beads were washed in 50 mM Tris-HCl pH 8 and subjected to on-bead digestion (below), and the resulting peptides were analyzed by LC–MS/MS. For SDS-PAGE and immunoblotting, bound proteins were eluted in 1*×* Laemmli buffer at 95 *^◦^*C for 10 min (1 mg mitochondrial equivalent).

**Lat1 immunoprecipitation** Immunoprecipitation of the PDH E2 subunit Lat1 was performed from isolated mitochondria using purified anti-Lat1 antibody coupled to magnetic beads (Dynabeads M-270 Epoxy). Beads were conjugated as described previously [57]: 8 mg beads were equilibrated in 0.1 M sodium phosphate buffer pH 7.4 and conjugated overnight at 30 *^◦^*C with 80 *µ*l purified Lat1 antibody, then washed sequentially with 0.1 M sodium phosphate buffer pH 7.4, 100 mM glycine-HCl, 10 mM Tris-HCl pH 8.8, 100 mM triethylamine, PBS, and 0.5% Triton X-100, and stored in PBS with 0.02% NaN_3_ at 4 *^◦^*C. Per sample, 2 mg mitochondria were solubilized in lysis buffer (10 mM Tris-HCl pH 7.4, 150 mM NaCl, 1*×* Roche cOmplete with EDTA) supplemented with 1% digitonin for 30 min at 4 *^◦^*C and cleared by centrifugation (13,000*× g*, 10 min, 4 *^◦^*C). Cleared lysate was incubated with the Lat1-coupled beads for 2 h at 4 *^◦^*C. A Δ*lat1* strain was processed in parallel as a specificity control. Four biological replicates were prepared for each condition in two independent experimental sessions (replicates 1–2 and replicates 3–4, respectively). Beads were washed three times with lysis buffer supplemented with 0.05% digitonin and three times with detergent-free high-salt lysis buffer (10 mM Tris-HCl pH 7.4, 250 mM NaCl, 1*×* Roche cOmplete with EDTA), and bound proteins were eluted in 1*×* Laemmli buffer at 95 *^◦^*C for 10 min. One sixth of each eluate was retained for immunoblot analysis; the remainder was resolved by SDS-PAGE into a stacking gel, and the stacked protein bands were excised and subjected to in-gel digestion, with the resulting peptides analyzed by LC–MS/MS.

**Pda1–TurboID proximity labeling** Cells expressing Pda1–TurboID or the matrix-localized Floaty–TurboID control were grown in YPGal medium and supplemented with 50 µM (final concentration) biotin for 30 min to promote biotinylation. Labeling was performed in whole, intact yeast cells prior to mitochondrial isolation. Mitochondria were isolated and solubilized in 1% SDS (5 min, 50 *^◦^*C), followed by dilution into RIPA buffer (50 mM Tris-HCl pH 7.5, 150 mM NaCl, 1% NP-40, 1 mM EDTA, 1 mM EGTA, 0.1% SDS, 0.5% sodium deoxycholate, protease inhibitor cocktail) supplemented with Benzonase nuclease, and biotinylated proteins were captured on streptavidin-coupled magnetic beads (50 µl beads per sample) for 1 h at 4 *^◦^*C with rotation. Beads were washed stringently (3*×* with 1 ml RIPA buffer, followed by 3*×* with 1 ml TAP buffer (50 mM HEPES-KOH pH 8.0, 100 mM KCl, 10% (v/v) glycerol, 2 mM EDTA) to remove residual detergent, transferring to a fresh tube after the first wash in each series) to remove non-covalently bound material. For immunoblot analysis, 1/6 of the washed beads was eluted by boiling in 1*×* Laemmli sample buffer (95 *^◦^*C, 10 min) and analyzed by immunoblotting, including streptavidin– HRP detection of total biotinylation. For mass spectrometry, the remaining beads were snap-frozen in liquid nitrogen and subjected to on-bead digestion (below).

**On-bead digestion** Following the final stringent washes, beads were transferred to fresh tubes and buffer-exchanged at 4 *^◦^*C to remove detergent, salts, and protease inhibitors: twice with 500 µl wash buffer 1 (15 mM Tris-HCl pH 7.4, 40 mM KCl, 2 mM EDTA), twice with 500 µl wash buffer 2 (50 mM Tris-HCl pH 7.4), and once with 100 µl wash buffer 3 (50 mM Tris-HCl pH 8.0). Proteins were digested on-bead by addition of 80 µl trypsin buffer (50 mM Tris-HCl pH 8.0, 2 M urea, 1 mM DTT) containing 0.3 µg Trypsin/Lys-C (Promega V5071) for 30 min at 37 *^◦^*C and 500 rpm with a heated lid. Supernatants were collected, and beads were washed twice with 60 µl wash buffer 4 (50 mM Tris-HCl pH 8.0, 1 M urea), pooling all supernatants. Samples were reduced with 4 mM DTT (30 min, 37 *^◦^*C), alkylated with 10 mM iodoacetamide (30 min, 25 *^◦^*C in the dark), and digested to completion with a further 0.2 µg Trypsin/Lys-C overnight at 25 *^◦^*C. Digestion was stopped with formic acid (1.33% final; pH 2–3 confirmed on indicator paper). Peptides were desalted on home-made C18 stage tips [58], vacuum-dried, and stored at *−*80 *^◦^*C.

**LC–MS/MS** LC–MS/MS was performed on two instruments, corresponding to the two sets of experiments. The Abf2–Spot pulldown samples were analysed on an Ultimate 3000 RSLCnano system (Thermo Fisher Scientific): peptides were desalted on a C18 trap column and separated on a 25 cm analytical column (75 µm inner diameter, 1.6 µm C18; Odyssey-IonOpticks) over a 50 min gradient from 2% to 35% (v/v) acetonitrile in 0.1% formic acid, and the effluent was electrosprayed directly into a Q Exactive HF mass spectrometer (Thermo Fisher Scientific) operated in data-dependent mode. Survey full-scan MS spectra (*m/z* 375–1600) were acquired at a resolution of *R* = 60,000 (at *m/z* 400) with an AGC target of 3 *×* 10^6^; the ten most intense ions with charge states of 2–5 were sequentially isolated to a target value of 1 *×* 10^5^ and fragmented at a normalized collision energy of 27%. Additional parameters were: spray voltage, 1.5 kV; no sheath or auxiliary gas flow; heated capillary temperature, 250 *^◦^*C; and ion-selection threshold, 33,000 counts. The Lat1 immunoprecipitation and Pda1–TurboID samples were analysed as described previously [59] on a nano-LC system (Ultimate 3000 RSLC, Thermo Fisher Scientific) coupled to an Impact II high-resolution Q-TOF (Bruker Daltonics) via a CaptiveSpray nano-ESI source (Bruker Daltonics). Peptides were trapped on an Acclaim PepMap nano-trap column (C18, 100 Å, 100 µm *×* 2 cm) and separated on an Acclaim PepMap RSLC analytical column (C18, 100 Å, 75 µm *×* 50 cm; both Thermo Fisher Scientific) over a 90 min linear gradient of 4–45% (v/v) acetonitrile at 250 nl/min, with the column held at 50 *^◦^*C. MS1 spectra (*m/z* 200–2000) were acquired at 3 Hz, and the 18 most intense precursors were selected for MS/MS at an intensity-dependent rate of 4–16 Hz, with a dynamic exclusion of 0.5 min.

**Database searching** Raw files were processed with MaxQuant [60] and searched against the *Saccharomyces cerevisiae* reference proteome (UniProt UP000002311, strain ATCC 204508/S288c). Trypsin was set as the protease with up to two missed cleavages; carbamidomethylation of cysteine was a fixed modification, and N-terminal acetylation and methionine oxidation were variable modifications; phosphorylation of serine, threonine and tyrosine was additionally included as a variable modification for the Lat1 dataset. A false discovery rate of 1% was applied at both the peptide and protein level, with a minimum peptide length of seven residues. The Lat1 immunoprecipitation, Pda1–TurboID, and Abf2 pulldown datasets were searched independently, the former two with MaxQuant v2.4.9.0 and the Abf2 pulldown with v2.0.1.0. Protein abundance was quantified by label-free quantification (LFQ) [61] without match-between-runs for the Lat1 and Pda1–TurboID datasets, and by intensity-based absolute quantification (iBAQ) with match-between-runs enabled (matching window 0.7 min, alignment window 20 min) for the Abf2-Spot pulldown.

**Data analysis: Abf2-Spot interactome** The MaxQuant proteinGroups output (Database searching, above) was filtered to remove reverse hits, potential contaminants, and entries identified only by site, and protein groups were retained if supported by at least one unique peptide (471 of 484 groups). Protein abundance was taken as iBAQ intensity; values were log_2_-transformed, with zero values treated as missing, and remaining missing values were imputed per sample from a down-shifted normal distribution (down-shift 1.8 SD, width 0.3 SD) with a fixed random seed for reproducibility. Enrichment was computed for the native and the crosslinked purification separately, as the background-corrected log_2_ fold change between the Abf2–Spot (bait) and the untagged control sample. Proteins were annotated as mitochondrial using the high-confidence mitochondrial proteome [19] and, for display, grouped into the PDH complex, nucleoid / mtDNA-maintenance factors, and mitochondrial gene-expression machinery (mitoribosomal subunits, translation factors, mtRNA-metabolism factors, and mtDNA-encoded proteins); the analysis was restricted to mitochondrial proteins. As this was a single exploratory purification without biological replicates, no significance testing was applied: enrichment is reported as background-corrected log_2_ fold change, its specificity assessed by dependence on formaldehyde crosslinking, and the principal findings were independently confirmed by immunoblotting (Fig. 1B). Analyses were performed in Python using pandas, NumPy, and matplotlib.

**Data analysis: Lat1 interactome** From the proteinGroups output, entries flagged as reverse hits, potential contaminants, or identified only by site were removed. LFQ intensities were log_2_-transformed, with zero values treated as missing. Proteins were retained if quantified in at least three of four replicates in at least one condition. Remaining missing values were imputed from a per-sample down-shifted normal distribution (down-shift 1.8 SD, width 0.3 SD), following the standard approach for label-free interactome data [62]; a fixed random seed ensured reproducibility. Enrichment of each protein in Lat1 versus control purifications was assessed by a two-sided Welch’s *t*-test, and *p*-values were corrected for multiple testing using the Benjamini–Hochberg procedure. Proteins with log_2_ fold change *>* 1 and adjusted *p*-value (*q*) *<* 0.05 were considered enriched. Enriched interactors were ranked by *π*-score, defined as log_2_ fold change *× −*log_10_*q* [18]; this ranking is descriptive and no category-level enrichment statistic was applied, as an affinity purification does not provide an unbiased proteome-wide background. Proteins were annotated as mitochondrial using the high-confidence mitochondrial proteome of Morgenstern *et al.* [19] and assigned to functional groups (PDH complex; nucleoid / mtDNA maintenance; large and small mitoribosomal subunits; mitochondrial translation factors; mtDNA-encoded proteins) using manually curated gene sets based on SGD annotation. Analyses were performed in Python using pandas, NumPy, SciPy, and matplotlib.

**Data analysis: Pda1-proximal proteome** From the proteinGroups output, contaminant, reverse-decoy, and only-identified-by-site entries were removed using the corresponding MaxQuant flag columns. LFQ intensities were log_2_-transformed, with zero values treated as missing. Proteins were retained if quantified in at least two of three replicates of the Pda1–TurboID group, and remaining missing values were imputed from a per-sample down-shifted normal distribution (down-shift 1.8 SD, width 0.3 SD), following the standard approach for label-free interactome data [62]; a fixed random seed ensured reproducibility. Imputation provides values for on/off comparisons, in particular against the untagged control, in which *∼*93% of values were missing owing to the absence of biotinylation. Differential enrichment was assessed by a two-sided Welch’s *t*-test on log_2_ LFQ intensities, with Benjamini–Hochberg (BH) correction for multiple testing. Proteins were called enriched at BH *q <* 0.05 and log_2_ fold change *>* 1 (2-fold). The Pda1-proximal proteome (foreground) was defined as the proteins meeting these criteria in both the Pda1–TurboID versus Floaty–TurboID contrast (spatial specificity) and the Pda1–TurboID versus untagged contrast (biotinylation dependence). The background for enrichment testing was the union of all proteins detected across the three conditions (*n* = 420), rather than the whole *S. cerevisiae* or published mitochondrial proteome, to restrict the test to specificity within the sampled compartment.

Curated gene sets representing mtDNA-associated machineries were derived from *S. cerevisiae* Gene Ontology annotations in the *Saccharomyces* Genome Database (via YeastMine): the mitochondrial large (GO:0005762; 47 genes, 36 detected) and small (GO:0005763; 36 genes, 28 detected) ribosomal subunits; the complete set of mitochondrial translation components (104 genes, 80 detected); a non-ribosomal translation set comprising translation factors and aminoacyl-tRNA synthetases, obtained by subtracting the ribosomal subunits from the complete translation set (22 genes, 16 detected); the mtDNA nucleoid / genome-maintenance set (24 genes, 21 detected); and, as comparison sets, TCA cycle enzymes (28 genes, 17 detected) and oxidative phosphorylation subunits (51 genes, 21 detected; SGD GO:0098803 and GO:0045259, manually curated to exclude the transcription factor Hap4 and the intron-encoded maturases Ai3/Ai4/Ai5*α*, which are not structural subunits). For each set, only genes detected in the experiment were included in the test, so the enrichment reflects specificity within the observable compartment. Over-representation among the foreground was assessed by hypergeometric testing against the detected background, with BH correction across sets. Only the large and small mitochondrial ribosomal subunits were significantly enriched; the complete translation set was also enriched, but this reflected inclusion of the ribosomal proteins, as the non-ribosomal translation set was not enriched (*q ≈* 0.11). To confirm that the ribosome enrichment was not an artifact of imputation, the analysis was repeated requiring detection in all three Pda1–TurboID replicates, such that no foreground protein carried imputed values; the large and small subunit enrichments were preserved (*q* = 9.5 *×* 10*^−^*^4^ and 2.0 *×* 10*^−^*^3^, respectively). Analyses were performed in Python using pandas, NumPy, and SciPy.

### SDS–PAGE and immunoblotting

Proteins were separated on 12%, 14% or 16% SDS–polyacrylamide gels and transferred to PVDF (2 mA/cm^2^ for 90 min) membranes. Membranes were blocked in TBS containing 5% (w/v) milk powder and probed with the primary antibodies at the indicated dilutions, followed by HRP-conjugated secondary antibodies listed in Supplementary Table S3. Signals were detected by enhanced chemiluminescence on a Vilber Fusion FX imaging system. Antibody specificity was validated against the corresponding deletion strains (Supplementary Fig. S4D).

### Digitonin solubilization and supernatant/pellet fractionation

To test the nucleic acid dependence of PDHc subunit solubility, freshly isolated mitochondria (600 µg in SEM buffer [250 mM sucrose, 1 mM EDTA, 10 mM MOPS-KOH, pH 7.2]) from wild-type, Δ*pda1*, Δ*pdb1*, or Δ*lat1* cells grown in YPD, YPG, or YPGal medium were divided into 100-µg aliquots and lysed in 50 µl lysis buffer (10 mM Tris-HCl pH 7.4, 150 mM NaCl, 1*×* cOmplete EDTA-free protease inhibitor cocktail, 1% [w/v] digitonin; protease inhibitor and digitonin added fresh) in the presence or absence of nuclease. For RNase treatment, 1 µl RNase I (Thermo Fisher) was added per sample. For DNase treatment, the lysis buffer was additionally supplemented with 5 mM CaCl_2_and 5 mM MgCl_2_, and 1 µl DNase I (Thermo Fisher) was added per sample. Samples were lysed for 30 min at 4 *^◦^*C. An aliquot corresponding to 50 µg protein (Total) was set aside and subjected to the same centrifugation step as the remaining material; the latter was cleared by centrifugation at 13,000 *×* g for 10 min at 4 *^◦^*C to separate soluble (Supernatant) from insoluble (Pellet) material. Pellets were resuspended in an equal volume of lysis buffer, such that Total, Supernatant, and Pellet fractions each represented an equivalent proportion of the starting material. All fractions were denatured in 1*×* Laemmli sample buffer (95 *^◦^*C, 10 min) and analyzed by SDS-PAGE and immunoblotting.

Nuclease activity was verified in parallel using purified nucleic acid substrates. For RNase controls, 1 µg of mitochondrial RNA was incubated in water, lysis buffer, or lysis buffer supplemented with RNase I. For DNase controls, 2 µl of purified plasmid DNA (362 ng/µl) was incubated in water, lysis buffer, or lysis buffer supplemented with DNase I (plus 5 mM CaCl_2_/MgCl_2_). Control samples were kept on ice for 30 min alongside the mitochondrial samples and then resolved by agarose gel electrophoresis.

For SDS-PAGE, each sample (33.3 µl total: 25 µl sample + 8.3 µl 4*×* Laemmli buffer) was loaded in duplicate onto two identical gels (16.65 µl per gel) to allow probing with multiple antibodies; loading volumes were adjusted proportionally to correct for pipetting error and ensure equal loading across samples. Gels were run at 90 V for 10 min, followed by 130 V for 75–90 min, and proteins were transferred at 2 mA/cm^2^ membrane for 90 min. Membranes were blocked for 1 h at room temperature. Anti-Pda1 primary antibody was incubated for 1 h at room temperature; anti-Pdb1, anti-Lat1, anti-Lpd1, and anti-Pdx1 primary antibodies were incubated overnight at 4 *^◦^*C. Secondary antibodies were incubated for 1 h at room temperature.

Equivalent fractions were analyzed by immunoblotting, and the fraction of each subunit in the pellet was quantified using Fiji (Fig. 3B).

### Sucrose density gradient centrifugation

Isolated mitochondria (7 mg) were thawed, pelleted, and resuspended in 3,500 µl lysis buffer (20 mM Tris-HCl pH 7.4, 0.5 M sucrose, 50 mM NaCl, 2 mM EDTA, 7 mM *β*-mercaptoethanol, 0.5% NP-40, and 1*×* cOmplete EDTA-free protease inhibitor cocktail). The lysate was split into two 1,750-µl aliquots and incubated with or without 1 µl RNase I (Thermo Fisher) for 5 min on ice; RNase activity was confirmed in parallel using mitochondrial RNA incubated in lysis buffer with or without RNase I. An aliquot (250 µl) of each lysate was reserved as input and diluted 1:5 in lysis buffer (150 µl lysate + 600 µl lysis buffer). A 1.5 ml volume of each lysate (*±* RNase) was loaded onto an 11-step discontinuous sucrose gradient (20–65% [w/v], 4.5% increments, 5 ml per step) prepared in gradient buffer (20 mM Tris-HCl pH 7.4, 50 mM NaCl, 2 mM EDTA, 7 mM *β*-mercaptoethanol, and 1*×* cOmplete EDTA-free protease inhibitor cocktail). Gradients were centrifuged at 110,000 *×* g (29,500 rpm in a swinging-bucket rotor) for 70 min at 4 *^◦^*C as described previously [12]. Seventeen 750-µl fractions were collected sequentially from the top of each gradient. From each fraction, 300 µl was reserved for nucleic acid extraction and 90 µl for protein analysis; the protein aliquot was mixed with 30 µl 4*×* Laemmli sample buffer, and 25 µl was loaded per lane on duplicate gels. Proteins were analyzed by SDS-PAGE and immunoblotting using antibodies against the PDHc subunits Pda1, Pdb1, Lat1, and Lpd1, and against Pdx1.

### Fluorescence microscopy

Cells were grown to mid-log phase in YPD medium (2% glucose) at 30°C, immobilized on ConA-coated Ibidi 8-well µ-slides, and imaged at 30°C on a Nikon Ti2-Eclipse wide-field fluorescence microscope equipped with a CFI Apochromat TIRF 100×/1.49 NA oil objective and a Photometrics Prime 95B 25 mm camera. Z-stacks were acquired to a total depth of 8 µm, with a step size of 0.2 µm per slice. Specific filter and acquisition set-ups can be made available upon request.

**DAPI-staining** For mtDNA staining, 2 *×* 10^6^ cells from mid-log cultures were harvested by centrifugation (3,000*× g*, 3 min) and resuspended in 1 ml SC medium containing 4 *µ*g/ml DAPI (4*^′^*,6-diamidino-2-phenylindole dihydrochloride). Cells were incubated for 15 min at 30 *^◦^*C with shaking (550 rpm) in the dark, washed once in PBS, and immobilized on ConA-coated Ibidi 8-well µ-slides as described above. Cells were overlaid with SC medium and imaged as above.

### Image quantification

Fluorescent channels were deconvolved using Huygens Essentials software (Scientific Volume Imaging).

**Pda1-spot analysis** Individual Pda1-NeonGreen puncta were segmented in 3D from the raw Z-stacks using a custom Fiji macro, which returned per-spot morphology and intensity measurement tables (volume, mean intensity, integrated density, min/max) for every cell, which were detected and segmented by Yeastmate [63]. Spots were classified per cell as “bright” or “dim” using a threshold of twice the median spot intensity of that cell, and the number of bright and dim spots per cell (and the corresponding fraction of the total) was quantified. Values from three biological replicates were pooled and visualized as superplots, with single-spot values overlaid by per-replicate means and the overall mean *±* SD.

**Pda1 spot Manders’ assay** The su9-mKate-labeled mitochondrial network of each cell was skeletonized in 3D using MitoGraph (0.11 *×* 0.11 *×* 0.2 µm xy/z voxel size) [64], yielding a set of network-node coordinates per cell. At each node, the local intensity of both the matrix marker and Pda1-NeonGreen was sampled from a 3 *×* 3 *×* 3-voxel neighbourhood in the raw image, restricting the analysis to positions within the mitochondrial network. Both intensity traces were thresholded per cell (Li’s method), and Manders’ colocalization coefficient (M1: fraction of matrix-marker signal residing in Pda1-positive voxels) was calculated for each cell, together with the Pearson correlation coefficient of the raw traces as a secondary measure. Per-cell coefficients were grouped by genotype (*ρ*^+^ vs. *ρ*^0^) and biological replicate, compared by *t*-test on replicate means, and displayed as superplots.

**Pda1-NG CAP treatment** The same MitoGraph-based network-sampling approach was used to extract matched matrix-marker and Pda1-NG intensities along the mitochondrial network for each cell across a chloramphenicol treatment timelapse. To prevent intensity drift over time (e.g., CAP-induced dimming or photobleaching) from confounding genuine changes in clustering, a single Pda1 intensity threshold was defined per cell from its own pre-treatment (*T*_0_) frame and applied to all later timepoints of that same cell (fixed-threshold Manders coefficient, M2); the matrix-marker signal was used unthresholded, since sampling points were already restricted to the network by construction. Manders coefficients at each timepoint were compared to the pre-treatment (*T*_0_) value using unpaired two-tailed Welch’s *t*-tests (unequal variance).

### Quantification of mtDNA copy number

Cells were harvested at mid-log phase from back-diluted overnight cultures by centrifugation (3.000 *× g*, 3 min, room temperature). Pellets were washed with sterile deionized water and stored at *−*80 *^◦^*C. Genomic DNA (gDNA) was extracted by bead-beating phenol-chloroform extraction as described previously [65]. DNA concentration was measured on a NanoPhotometer (Implen N60-Touch), and all samples were diluted to 0.1 ng/µl. Quantitative PCR targeting *ACT1* and *COX1* was performed as described previously [4]. Mitochondrial DNA (mtDNA) abundance was calculated as the ratio of *COX1* to *ACT1*, and relative abundance was determined by normalizing the mean of biological replicates for each sample strain to that of the reference strain.

### Petite frequency assay

To determine petite frequency, approximately 200 cells from mid-log-phase cultures grown at the indicated temperature and for the indicated duration were plated onto YPG plates supplemented with 0.1% glucose. Colonies were scored after 2–3 days at 30 *^◦^C* as either large (respiratory-competent, rho^+^) or small (petite, rho^0^/rho^-^) using a semi-automated image-analysis pipeline combining Fiji [66] and a custom Python script. Petite frequency was expressed as the percentage of petite colonies relative to the total number of colonies scored.

### Mild CAP treatment for petite analysis

Cells were grown overnight and back-diluted the following morning to an OD_600_ of 0.1. Chloram-phenicol was added to YPD medium at a final concentration of 0.5 mg/ml, and cells were incubated for 5 h at 30 *^◦^*C. Petite frequency was then assessed as described above.

### Monitoring mitochondrial translation via Cox2-sfGFP reporter

Monitoring of mitochondrial translation via the mitochondrially encoded superfolder GFP reporter (*sfGFP*^m^, integrated at the *cox2* locus) was performed as described previously [33]. Cells were grown overnight at 30 *^◦^*C in YPD then back-diluted the following morning into YPG. For the temperature-shift condition, cultures were grown for 11 h at either 30 *^◦^*C or 37 *^◦^*C. For chloramphenicol (CAP) treatment, cultures were grown at 30 *^◦^*C and treated with 1 mg/ml CAP for 5 h. In all cases, cultures were maintained in mid-log phase until sampling. Samples were analysed on a BD Accuri C6 Plus flow cytometer. Initial gating was performed with floreada.io and subsequent analysis with custom Python scripts. To exclude non-cell events (e.g. debris or agar particles), cells were gated on forward-scatter area versus forward-scatter height (FSC-A vs. FSC-H).

### Tetrad analysis

Haploid cells of opposing mating types grown to mid-log phase were mixed in a 1:1 ratio, pelleted (3 min, 3,000 *×* g), resuspended in 75 µl YPD, and plated for mating on YPD plates for 2.5 h at 30 *^◦^*C. Diploid cells were selected by picking approximately 30 morphologically characteristic zygotes per mating using a SporePlay+ dissection microscope (Singer Instruments). To confirm the diploid state of the resulting colonies, each colony was plated on medium containing both antibiotics, selecting for the resistance markers of the two mating partners. Confirmed diploids were streaked onto sporulation plates (3 g/L potassium acetate, 0.48 g/L uracil dropout supplement [D9535], 12.5 mg/L uracil, 2% agar, adjusted to pH 5.5) and incubated for 5–7 days at 25 *^◦^*C. For tetrad dissection, sporulated cells were pelleted and resuspended in 20 µl of 5 mg/ml zymolyase 20T (Amsbio, 120491-1) in 1.2 M sorbitol, incubated for 10 min at 25 *^◦^*C, and diluted with 180 µl of 1.2 M sorbitol to digest the ascus wall. Spores were dissected using the SporePlay+ microscope and incubated for 48 h at 30 *^◦^*C prior to imaging with a Vilber Fusion FX imaging system. Resulting tetrads were analyzed for 2:2 segregation of mating type (MAT*a* or MAT*α*) and for segregation of antibiotic resistance cassettes.

### RNA-Seq

**RNA isolation** Total RNA was isolated from wild-type and Δ*pda1* cells (three biological replicates per genotype) using a protocol adapted from Schrott & Osman [4]. For each sample, 30 OD_660_ units of mid-log-phase cells were harvested (3,146 *×* g, 5 min, room temperature), washed once in ice-cold water, snap-frozen, and stored at *−*80 *^◦^*C. All subsequent steps were carried out on ice using nuclease-free reagents. Cell pellets were resuspended in 700 µl TRIzol reagent (Invitrogen, 15596026) and disrupted with zirconium beads by vortexing (3 *×* 60 s at maximum speed, with cooling on ice between cycles). Lysates were cleared (12,000 *×* g, 5 min, 4 *^◦^*C), extracted with 140 µl chloroform, incubated for 10 min at room temperature, and centrifuged (12,000 *×* g, 10 min, 4 *^◦^*C). The aqueous phase was re-extracted with an equal volume of phenol/chloroform/isoamyl alcohol, and RNA was precipitated with an equal volume of isopropanol (10 min on ice; 12,000 *×* g, 10 min, 4 *^◦^*C). Pellets were washed with 75% ethanol (prepared in nuclease-free water), air-dried, and resuspended in 50 µl nuclease-free water. Residual DNA was removed by treating 20 µg of RNA with Turbo DNase (Thermo Fisher,AM2238) for 30 min at 37 *^◦^*C, followed by clean-up on a Monarch RNA purification kit (New England Biolabs) with elution in 30 µl nuclease-free water. RNA concentration was determined by NanoDrop spectrophotometry.

**RNA-Sequencing** Ribosomal RNA was depleted from total RNA, and strand-specific libraries were constructed by the dUTP method: following RNA fragmentation, first-strand cDNA was synthesized with random hexamer primers, and dUTP was incorporated in place of dTTP during second-strand synthesis, after which libraries were completed by end repair, A-tailing, adapter ligation, size selection, USER enzyme digestion, and PCR amplification. Libraries were quantified by Qubit and qPCR, sized on a Bioanalyzer, and sequenced on an Illumina instrument in paired-end mode (2*×*150 bp). Raw reads were quality-filtered by the provider to remove adapter-containing reads, reads with *>*10% ambiguous bases, and reads with *>*50% low-quality (*Q ≤* 5) bases; all libraries had a Q20 base fraction above 97.9%.

**RNA-Seq data analysis** Downstream analysis was performed on the Galaxy platform [67]. Clean paired-end reads were aligned with RNA STAR [68] to a custom *S. cerevisiae* reference genome in which the S288c mitochondrial sequence (reference assembly R64, NCBI GCF_000146045.2) was replaced by the assembled mitochondrial genome of the LacO-tagged strain (yCO380), and per-gene read counts were generated with STAR’s --quantMode GeneCounts option using a matching gene-model annotation carrying the manually curated LacO-mtDNA features. One wild-type replicate was excluded prior to differential-expression analysis, as it segregated as a clear outlier from all other samples by principal-component analysis; the remaining two wild-type and three Δ*pda1* replicates were used for the analysis. Differential expression between Δ*pda1* and wild-type was determined with DESeq2 v1.40.2 [69] using a single-factor design (genotype), with the Wald test and Benjamini–Hochberg correction, using default parameters (median-of-ratios normalization, parametric dispersion fit). Transcripts with an adjusted *p*-value *<* 0.1 were considered differentially expressed. Analysis was run under R 4.3.1.

### Statistical analysis

Analyses were performed in Python (SciPy). Data are presented as mean *±* SD from *n* independent biological replicates, with *n* given in the corresponding figure legends. Pairwise comparisons of petite frequency and mtDNA copy number used unpaired two-tailed Welch’s *t*-tests (unequal variance), and Cox2–sfGFP reporter comparisons used paired two-tailed *t*-tests on matched per-replicate median intensities. In all cases, each strain was compared to the corresponding wild-type or wild-type-rescue control using pre-specified pairwise comparisons; as these were planned a priori against a common reference, *P* -values are reported without correction for multiple comparisons. Significance is indicated as *∗ p <* 0.05, *∗∗ p <* 0.01, *∗ ∗ ∗ p <* 0.001; n.s., not significant. Statistical analyses for the mass-spectrometry, image analysis and RNA-sequencing data are described in the corresponding Methods sections.

### Data and code availability

Mass spectrometry proteomics data will be deposited in the PRIDE database. All custom analysis scripts and Fiji macros used in this study are publicly available in the GitHub repository https://github.com/osman-mtDNA-lab/PDHc_Manuscript.

## Acknowledgments

We are grateful to the following colleagues for kindly providing antibodies: Prof. Dr. Dejana Mokranjac (Tim17, Tim23, Tim44, Tom40, Yme1) and Prof. Dr. Chris Meisinger (Pda1). We thank Tanja Kautzleben for assistance with media and plate preparation and for general laboratory management, and Sylvia Berngehrer, Tatiana Schubert, and Sonja Kristo for glassware washing and autoclaving services. We particularly thank the “Mito-Club” and the Mokranjac lab for stimulating discussions. This work was supported by a European Research Council grant (ERCStG-714739 IlluMitoDNA), a DFG grant (OS 410/3-1), and a Human Frontier Science Program grant (RGP021/2023) awarded to C.O..

## Author contributions

Johannes Hagen (Conceptualization [equal], Data curation [equal], Formal analysis [equal], Investigation [equal], Methodology [equal], Validation [lead], Visualization [lead], Writing—original draft [equal], Writing—review & editing [equal]), Nupur Sharma (Conceptualization [equal], Data curation [equal], Formal analysis [equal], Investigation [equal], Methodology [equal], Validation [supporting], Visualization [supporting]), Felix Thoma (Investigation [supporting], Formal analysis [supporting], Methodology [supporting]), Veronika Iskra (Investigation [supporting], Formal analysis [supporting]), Serena Schwenkert (Data curation [supporting], Formal analysis [supporting], Investigation [supporting], Methodology [supporting]), Simon Schrott (Formal analysis [supporting], Investigation [supporting], Methodology [supporting], Visualization [supporting]), Ignasi Forné (Data curation [supporting], Formal analysis [supporting], Investigation [supporting], Methodology [supporting]), Julia Weisenseel (Investigation [supporting]), Mengqiao Yang (Investigation [supporting]), Nadja Lebedeva (Investigation [supporting]), and Christof Osman (Conceptualization [equal], Funding acquisition [lead], Project administration [lead], Resources [lead], Supervision [lead], Writing—original draft [equal], Writing—review & editing [equal]).

