## Supplementary Figures for "RNA-dependent association of the pyruvate dehydrogenase complex with mtDNA-containing assemblies supports mitochondrial translation"

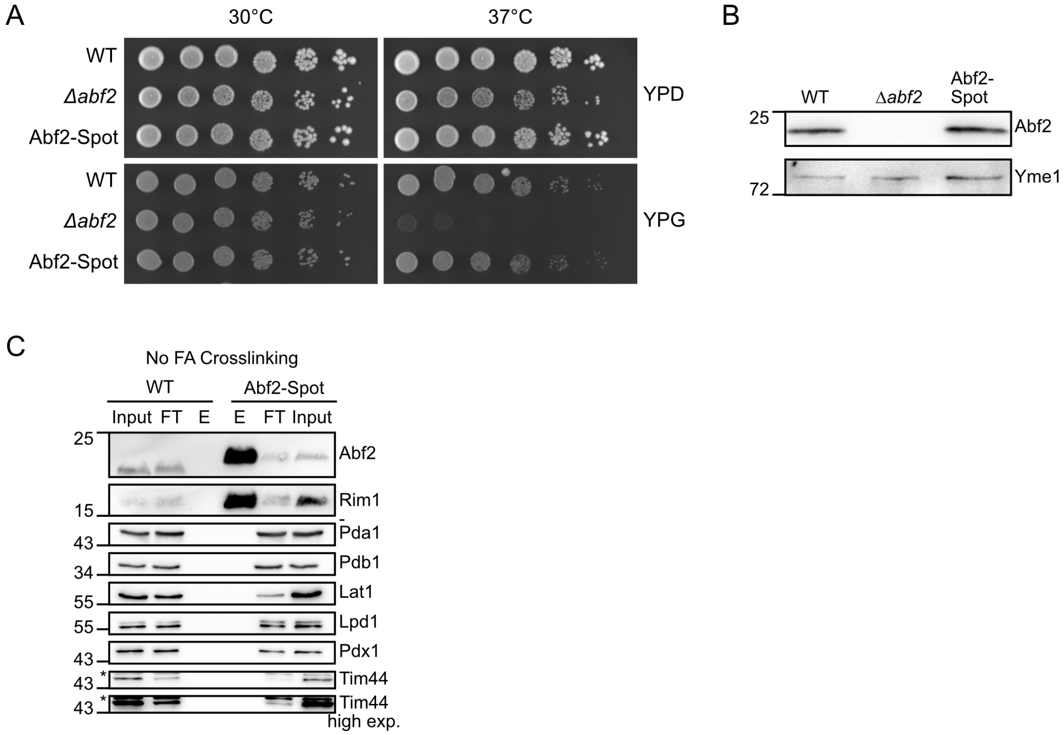


Figure S1: **Abf2–Spot is functional and co-purifies PDH subunits only after crosslinking. (A)** Growth analysis of wild-type (WT), Δ*abf2*, and Abf2–Spot cells on YPD and YPG at 30 °C and 37 °C. **(B)** Immunoblot of whole-cell extracts from WT, Δ*abf2*, and Abf2–Spot cells probed for Abf2 (Yme1, control). Molecular masses (kDa) are indicated at left. **(C)** Immunoblot of Abf2–Spot affinity purifications performed *without* formaldehyde crosslinking, comparing WT and Abf2–Spot mitochondria. Input, flow-through (FT), and eluate (E) fractions were probed with the indicated antibodies (Tim44 – higher-exposure below – control; asterisk, non-specific band). Molecular masses (kDa) are indicated at left.

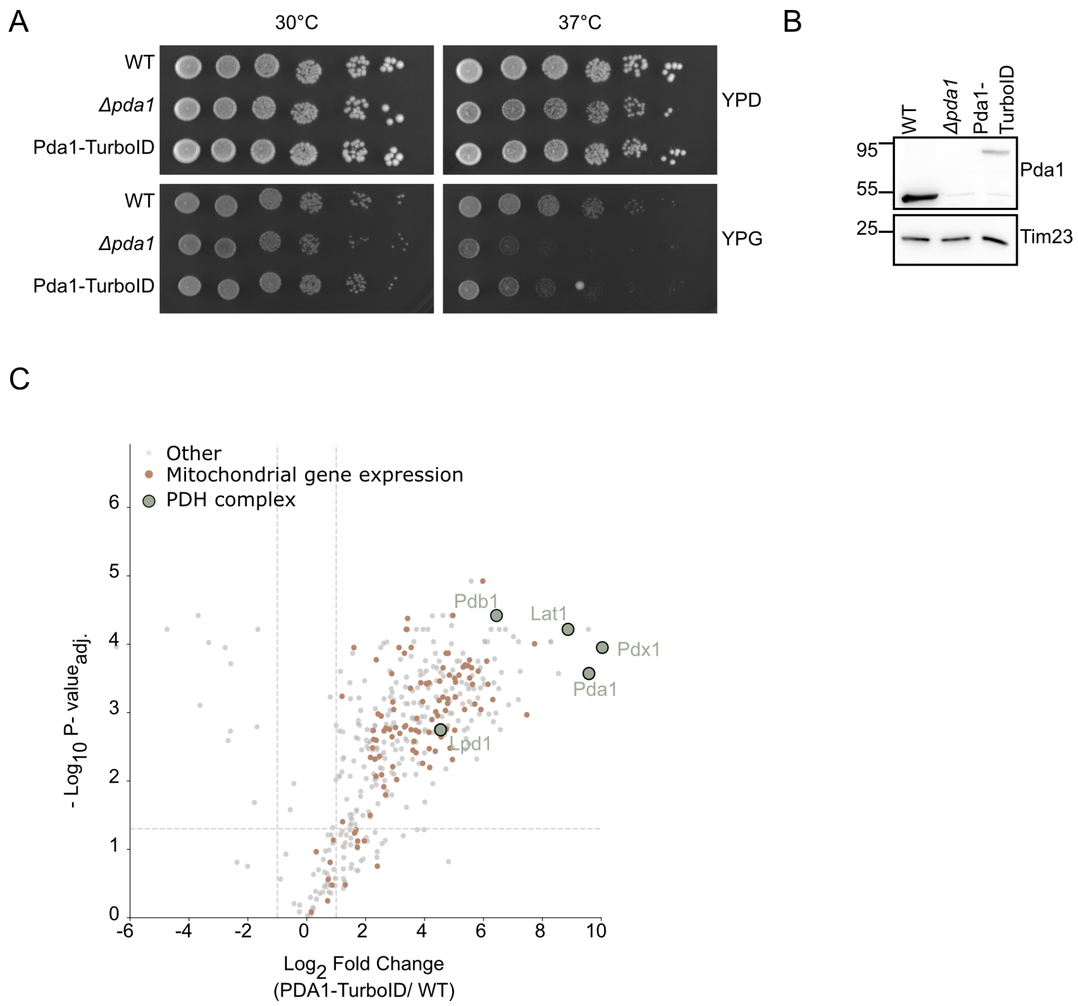


Figure S2: **Validation of the Pda1–TurboID fusion strain.** **(A)** Growth analysis of wild-type (WT), Δ*pda1*, and Pda1–TurboID cells on YPD and YPG at 30 °C and 37 °C. **(B)** Immunoblot of whole-cell extracts from WT, Δ*pda1*, and Pda1–TurboID cells probed for Pda1 (Tim23, control). Molecular masses (kDa) are indicated at left. **(C)** Volcano plot of streptavidin-purified proteins, Pda1–TurboID versus untagged wild-type control (Welch's *t*-test, Benjamini–Hochberg correction). PDH complex subunits (green) and mitochondrial gene-expression factors (orange) are highlighted; all other proteins are shown in grey. Dashed lines indicate the significance and fold-change thresholds ($q < 0.05$, $\log_2$ fold change $> 1$).

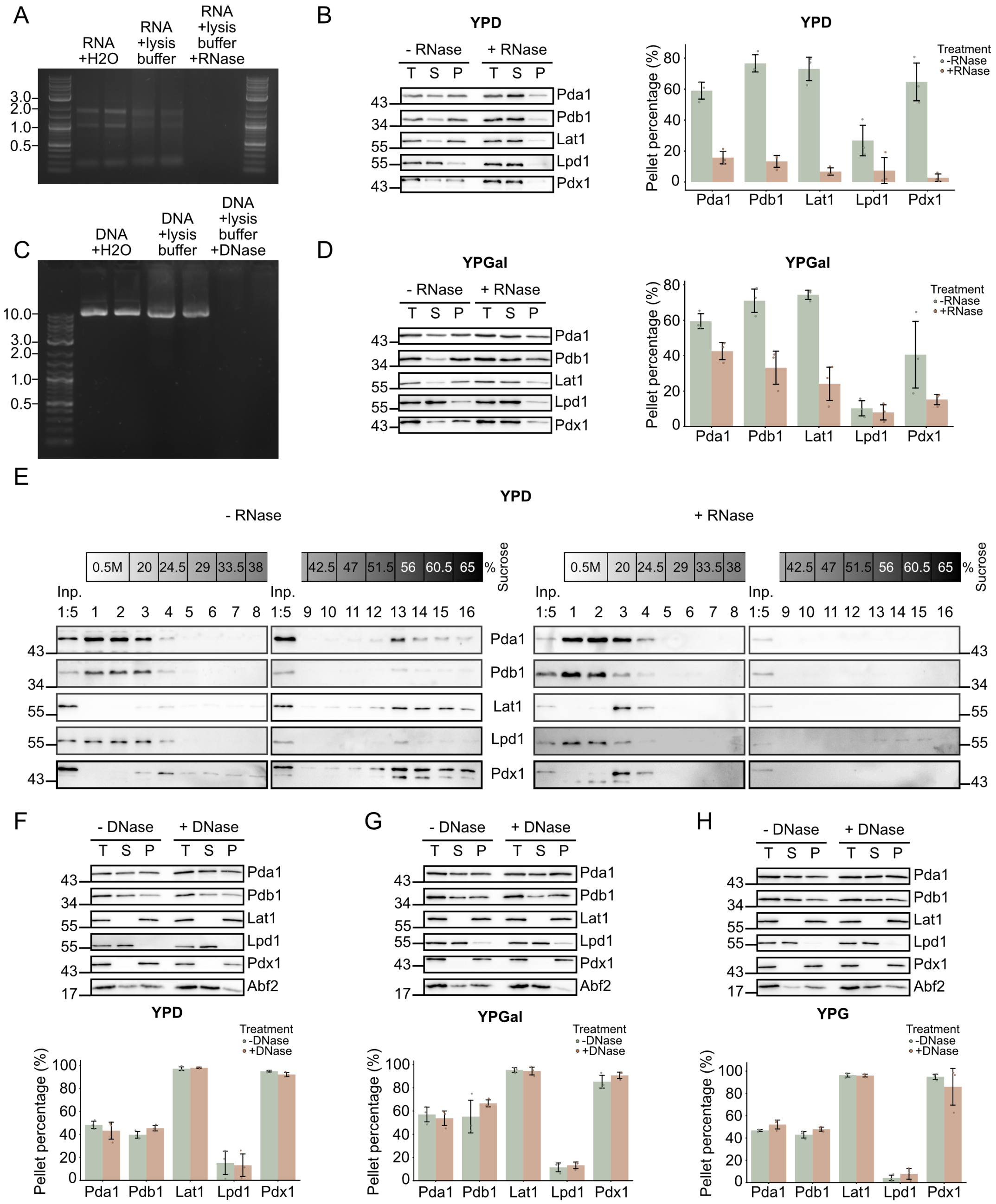


Figure S3: **The pellet association of PDH is RNA-dependent and DNA-independent across carbon sources.** **(A)** Agarose gel confirming RNase activity: purified mitochondrial RNA incubated in water, lysis buffer, or lysis buffer with RNase. DNA ladder sizes (kb) are indicated on the left. **(B)** Centrifugation-based fractionation of digitonin-solubilized mitochondria from YPD-grown cells into total (T), supernatant (S), and pellet (P) fractions, with or without RNase, analyzed by immunoblotting (left), with quantification of the pellet fraction (right). **(C)** Agarose gel confirming DNase activity: purified plasmid DNA incubated in water, lysis buffer, or lysis buffer with DNase. DNA ladder sizes (kb) are indicated on the left. **(D)** As in (B), for YPGal-grown cells. **(E)** Sucrose density gradient centrifugation (20–65%) of digitonin-solubilized mitochondria from YPD-grown cells, with or without RNase. Input and fractions 1–16 are shown. **(F–H)** Centrifugation-based fractionation as in (B), performed with or without DNase, for YPD (F), YPGal (G), and YPG (H) grown cells; immunoblots (top) and quantification of the pellet fraction (bottom). Abf2 is shown as a control. All quantifications: $n = 3$ biological replicates, mean ± SD.

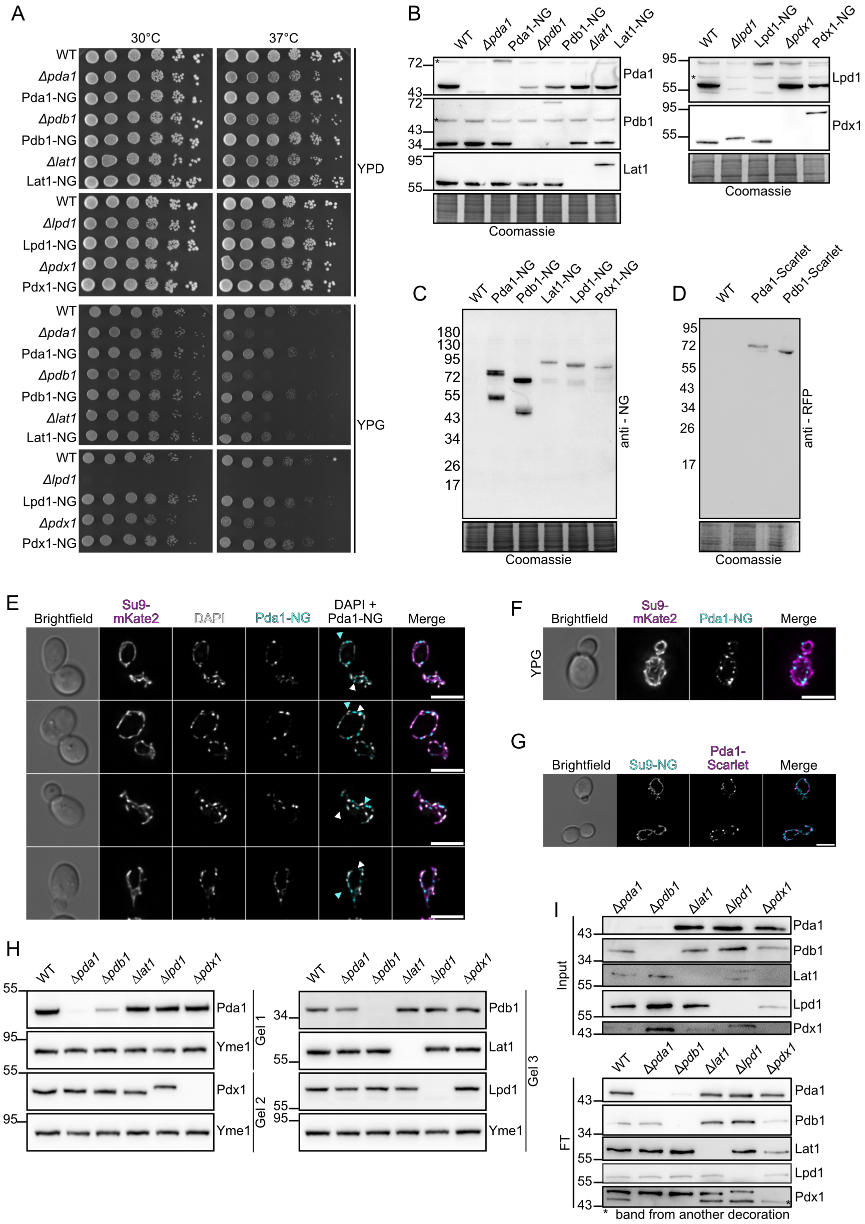


Figure S4: **Validation of fluorescently tagged PDH subunits and supporting analyses of Pda1 foci.** **(A)** Growth analysis of the indicated deletion strains and their mNeonGreen-tagged counterparts on YPD and YPG at 30 °C and 37 °C. **(B)** Immunoblots of the indicated PDH subunits in wild-type, deletion, and NG-tagged strains; Coomassie staining is shown as a loading control. Asterisks mark non-specific bands. **(C)** Anti-mNeonGreen (NG) immunoblot of cells expressing the indicated PDH subunit–NG fusions; Coomassie staining is shown as a loading control. All fusions are detected at their predicted full-length mass; a faster-migrating species (offset by ∼23 kDa) is present for Pda1, Pdb1, Lat1, and Lpd1. **(D)** Anti-RFP immunoblot of cells expressing Pda1–mScarlet or Pdb1–mScarlet; Coomassie staining is shown as a loading control. Each fusion is detected as a single full-length species. **(E)** Additional Pda1–NG cells co-stained with DAPI, corresponding to Fig. 4F. Cyan arrowheads, Pda1–NG foci lacking a coincident DAPI signal; white arrowheads, DAPI-stained nucleoids with an adjacent Pda1–NG focus. Scale bars, 5 µm. **(F)** Fluorescence microscopy of Pda1–NG and Su9–mKate2 in YPG-grown cells. Scale bar, 5 µm. **(G)**Fluorescence microscopy of cells co-expressing Pda1–mScarlet and the matrix marker Su9–NG. Scale bar, 5 µm. **(H)** Immunoblots of the indicated PDH subunits in wild-type and PDH deletion strains (Yme1, control). **(I)** Immunoblots of input and flow-through (FT) fractions from the Lat1 immunoprecipitations shown in Fig. 4D, probed for the indicated PDH subunits. Asterisks mark bands from another decoration.

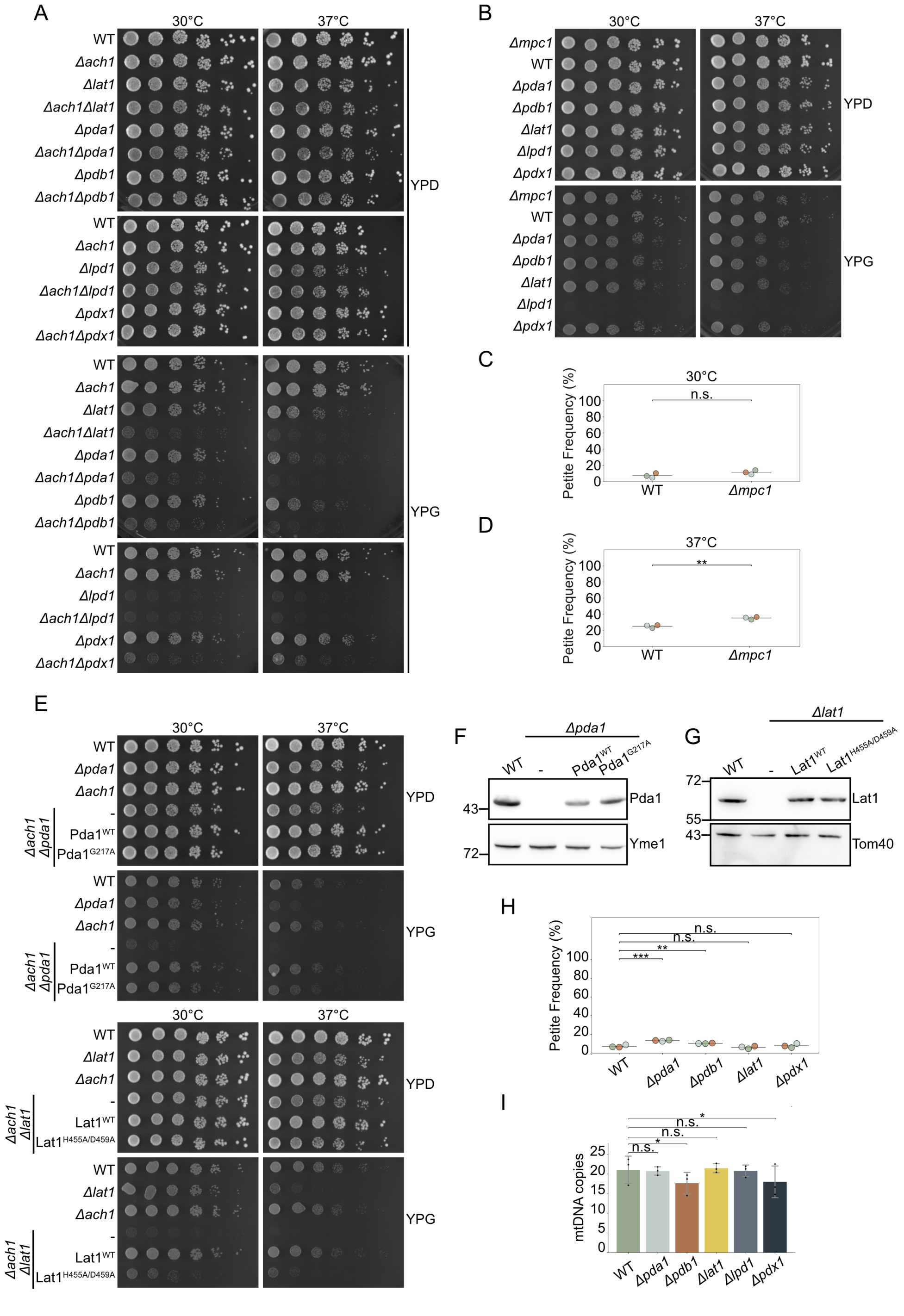


Figure S5: **Genetic interactions with *ACH1*, expression of catalytically inactive variants, and mtDNA phenotypes at 30 °C.** **(A)** Growth analysis of the indicated single and Δ*ach1* double mutants on YPD and YPG at 30 °C and 37 °C. **(B)** Growth analysis of Δ*mpc1* alongside wild-type and PDH deletion strains on YPD and YPG at 30 °C and 37 °C. **(C, D)** Petite frequency of wild-type and Δ*mpc1* cells at 30 °C (C) and 37 °C (D). **(E)** Growth analysis of Δ*ach1* Δ*pda1* and Δ*ach1* Δ*lat1* cells expressing no variant (−), the wild-type protein, or the catalytically inactive variant ($Pda1^{G217A}$, $Lat1^{H455A/D459A}$) on YPD and YPG at 30 °C and 37 °C. **(F)** Immunoblot of Pda1 in Δ*pda1* cells expressing no variant (−), $Pda1^{WT}$, or $Pda1^{G217A}$ (Yme1, control). Molecular masses (kDa) are indicated at left. **(G)** Immunoblot of Lat1 in Δ*lat1* cells expressing no variant (−), $Lat1^{WT}$, or $Lat1^{H455A/D459A}$ (Tom40, control). Molecular masses (kDa) are indicated at left. **(H)** Petite frequency of wild-type and the indicated PDH deletion strains at 30 °C. **(I)** Mitochondrial DNA copy number of wild-type and the indicated PDH deletion strains at 30 °C, determined by qPCR (*COX1*/*ACT1*). All quantifications: $n = 3$ biological replicates, mean ± SD.

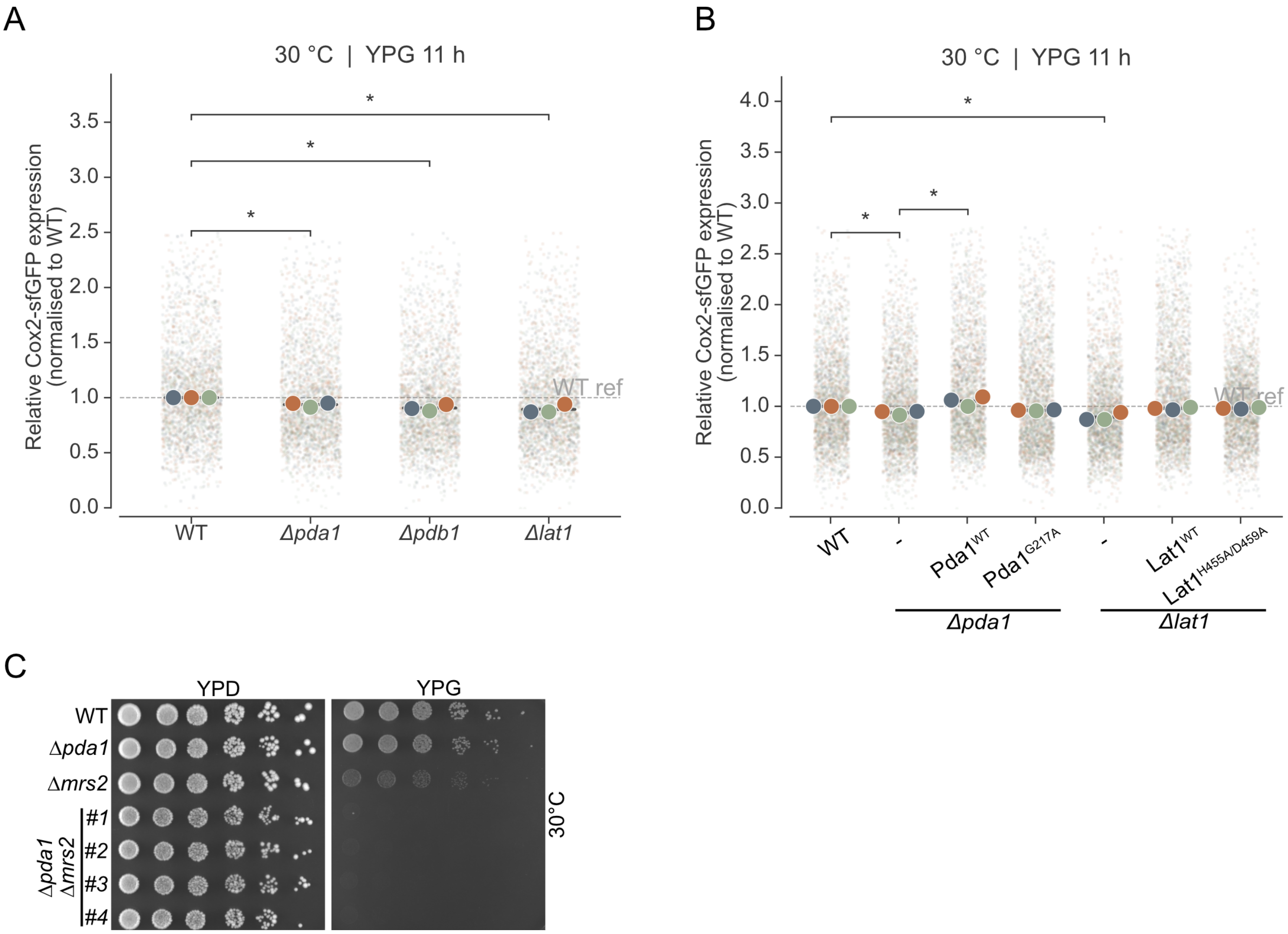


Figure S6: **Mitochondrial translation reporter output at 30 °C and genetic interaction of *PDA1* with *MRS2* .** **(A)** Cox2–sfGFP reporter expression in wild-type, Δ*pda1*, Δ*pdb1*, and Δ*lat1* cells grown in YPG for 11 h at 30 °C, normalized to wild-type (WT ref, dashed line). $n = 3$ biological replicates. **(B)**Cox2–sfGFP reporter expression in Δ*pda1* and Δ*lat1* cells expressing no variant (−), the wild-type protein, or the catalytically inactive variant ($\text{Pda1}^{\text{G217A}}$, $\text{Lat1}^{\text{H455A/D459A}}$), grown as in (B) and normalized to wild-type. Plotted as in (B). $n = 3$ biological replicates. **(C)** Growth analysis of wild-type, Δ*pda1*, Δ*mrs2*, and four independent Δ*pda1* Δ*mrs2* double-mutant segregants (#1–4), each from a separate tetrad, on YPD and YPG at 30 °C. Cox2–sfGFP reporter expression in Δ*pda1* and Δ*lat1* cells expressing no variant (−), the wild-type protein, or the catalytically inactive variant ($\text{Pda1}^{\text{G217A}}$, $\text{Lat1}^{\text{H455A/D459A}}$), grown as in (B) and normalized to wild-type. Plotted as in (B). $n = 3$ biological replicates.
